# The Set1 complex is dimeric and acts with Jhd2 demethylation to convey symmetrical H3K4 trimethylation

**DOI:** 10.1101/477646

**Authors:** Rupam Choudhury, Sukhdeep Singh, Senthil Arumugam, Assen Roguev, A. Francis Stewart

## Abstract

Epigenetic modifications can maintain or alter the inherent symmetry of the nucleosome however the mechanisms that deposit and/or propagate symmetry or asymmetry are not understood. Here we report that yeast Set1C/COMPASS is dimeric and consequently symmetrically trimethylates histone 3 lysine 4 (H3K4me3) on promoter nucleosomes. Mutation of the dimer interface to make Set1C monomeric abolished H3K4me3 on most promoters. The most active promoters, particularly those involved in the oxidative phase of the yeast metabolic cycle, displayed H3K4me2, which is normally excluded from active promoters, and a subset of these also displayed H3K4me3. In wild-type yeast, deletion of the sole H3K4 demethylase, Jhd2, has no effect. However in monomeric Set1C yeast, Jhd2 deletion increased H3K4me3 levels on the H3K4me2 promoters. Notably, the association of Set1C with the elongating polymerase was not perturbed by monomerisation. These results imply that symmetrical H3K4 methylation is an embedded consequence of Set1C dimerism and that Jhd2 demethylates asymmetric H3K4me3. Consequently, rather than methylation and demethylation acting in opposition as logic would suggest, a dimeric methyltransferase and monomeric demethylase co-operate to eliminate asymmetry and focus symmetrical H3K4me3 onto selected nucleosomes. This presents a new paradigm for the establishment of epigenetic detail.

## INTRODUCTION

The fundamental unit of eukaryotic chromatin, the nucleosome, is composed of four histone pairs (H2A, H2B, H3, H4) symmetrically arranged around a dyad axis (Luger et al. 1997). Consequentially it has been naturally presumed that nucleosomes in chromatin are symmetrical. However, high resolution epigenetic mapping has identified asymmetric distributions of post-translational modifications and histone variants within nucleosomes near transcription start sites. In particular, bivalent nucleosomes with histone 3 lysine 27 (H3K27) bi- or tri-methylation (me2/me3) on one H3 tail and H3K4me3 or H3K36me3 on the other, as well as asymmetric distribution of H4K20me1, have been identified in mammalian cells (Voigt et al. 2012; Shema et al. 2016). In yeast, asymmetric distributions of H3K9ac and H2A.Z, indicate new subtleties in promoter architecture (Rhee et al. 2014; Ramachandran et al. 2015). Asymmetric nucleosomes clearly have the potential to contribute to the regulation of, or convey directional orientation within, chromatin. To date very few studies have explored the mechanisms that convey epigenetic symmetries or asymmetries. In the course of studies on the structure of the Set1 complex, Set1C, which is the sole H3K4 methyltransferase in *Saccharomyces cerevisiae*, we unexpectedly encountered this issue.

Amongst the circuitries involved in epigenetic regulation, H3K4 methylation is one of the most widely conserved. In all eukaryotes, H3K4me3 on promoter nucleosomes is a binding site for the major protein complexes involved in launching transcription from active promoters including TFIID (Vermeulen et al. 2007; 2010). Furthermore, the size of the H3K4me3 peak equates with the quantity of mRNA production (Howe et al. 2017; Soares et al. 2017). Nevertheless, the function of the H3K4 methylation system remains enigmatic and no unifying regulatory explanation has emerged (Howe et al. 2017; Woo et al. 2017). This is due to several conundrums including (i) its apparent dispensability for transcription in yeast (Margaritis et al. 2012; Lenstra et al. 2011; Weiner et al. 2012) and *Drosophila* (after somatic mutagenesis; (Hödl and Basler 2012), (ii) the counterintuitive finding that loss of H3K4 methylation in yeast provokes more increased than decreased mRNA expression (Margaritis et al. 2012; Weiner et al. 2012); and (iii) rather than central transcriptional functions, in higher eukaryotes the H3K4 methyltransferases are only required for very specialized developmental roles (Glaser et al. 2009; Bledau et al. 2014; Ernst et al. 2004; Lee et al. 2013). The identification of H3K4 demethylases has compounded the enigma. H3K4me2/3 can be removed by the Jarid1/Kdm5 class of Jumonji-domain demethylases (Liang et al. 2007; Klose et al. 2006; Huang et al. 2010). Deletion of the only *S.cerevisiae* H3K4 demethylase, Jhd2, has very little effect on gene expression (Ramakrishnan et al. 2016). Similarly, functional studies on the four mouse Jarid1 genes (Kdm5a-d) are confusing because ablation reveals incompletely penetrant and variegating phenotypes (Kidder et al. 2014; Schmitz et al. 2011; Catchpole et al. 2011; Scandaglia et al. 2017).

The eight subunit yeast Set1 complex, Set1C, was the first H3K4 methyltransferase complex to be biochemically defined (Roguev et al. 2001). It was also described as the seven subunit COMPASS (COMplex of Proteins Associated with Set1;(Miller et al. 2001), with the missing subunit and *in vitro* H3K4 enzymatic proof added later (Krogan et al. 2002). Set1C/COMPASS is centered on a four membered subcomplex composed of two heteromeric interactions between Swd1/Swd3 and Bre2/Sdc1 (Roguev et al. 2001; South et al. 2010; Tremblay et al. 2014; Avdic et al. 2011). This subcomplex is the highly conserved scaffold for all Set1/Trithorax-type H3K4 methyltransferases and is now termed WRAD (Ernst and Vakoc 2012) after the mammalian homologues WDR5, RBBP5, ASH2L, and DPY30 (corresponding to Swd3, Swd1, Bre2/Ash2 and Sdc1 in yeast).

Dpy30 was first identified as a protein required for dosage compensation in *C. elegans* (Hsu et al. 1995). When we found a Dpy30 homologue in Set1C (Roguev et al. 2001), we noted that Dpy30 homologues include a region similar to the dimerization interface of the regulatory subunit RIIa of protein kinase A, which is composed of two alpha helices that form a four-helix bundle during dimerization (Newlon et al. 1999). We proposed that yeast Set1C is dimeric with the Dpy30 homologue, Sdc1, as the dimer interface (Roguev et al. 2003; Dehé et al. 2006). Structural studies with peptides confirmed that the RIIa homology region in DPY30 dimerizes to form an X bundle nearly identical to the dimerization/docking (D/D) domain of PKA RIIa (Wang et al. 2009; Zhang et al. 2015a; Tremblay et al. 2014).

Despite these indications, the contribution of the Dpy30/Sdc1 dimerization interface to the structure and function of H3K4 methyltransferases has not been investigated, possibly because Dpy30 is not required for enzyme activity *in vitro* (Dou et al. 2005; Li et al. 2016) (although inclusion of Dpy30 stimulates activity(Jiang et al. 2011). Here we show that yeast Set1C is dimeric and dimerization relies on the Dpy30/Sdc1 dimer interface. Using alanine mutagenesis of the dimer interface, we observed dramatic changes in H3K4 methylation profiles in yeast strains expressing wild type dimeric or mutant monomeric Set1C. Insight into the H3K4 methylation profiles was further revealed by deletion of the H3K4 demethylase, Jhd2. These findings present a new paradigm for the mechanics of H3K4 methylation and demethylation in eukaryotic epigenetic regulation. Furthermore, our findings extend the recent insights acquired from the high resolution structures of the monomeric Set1C/COMPASS enzymatic core (Hsu et al., 2018; Qu et al., 2018).

## RESULTS

### Set1C dimerization is mediated by Sdc1

To evaluate whether Sdc1 contributes to multimerization of Set1C, we used size exclusion chromatography (SEC) to examine the complex with and without Sdc1. Wildtype (wt) Set1C is approximately 1000kDa (Miller et al. 2001; Roguev et al. 2003), however Set1C lacking Sdc1 is approximately 400kDa (Fig. 1A-C). Two discrete *sdc1* mutations, either to delete the Dpy-30 homology box (*sdc1^Δ121-165^*) or point mutate critical residues within it (*sdc1\**; V129A, L133A, L134A; Fig. 1A, Supplemental Fig. 1A), which were chosen according to previous dimerization mutations in RIIa (Newlon et al. 1999), also reduced the apparent size of Set1C to 400kDa.

**Figure 1.**
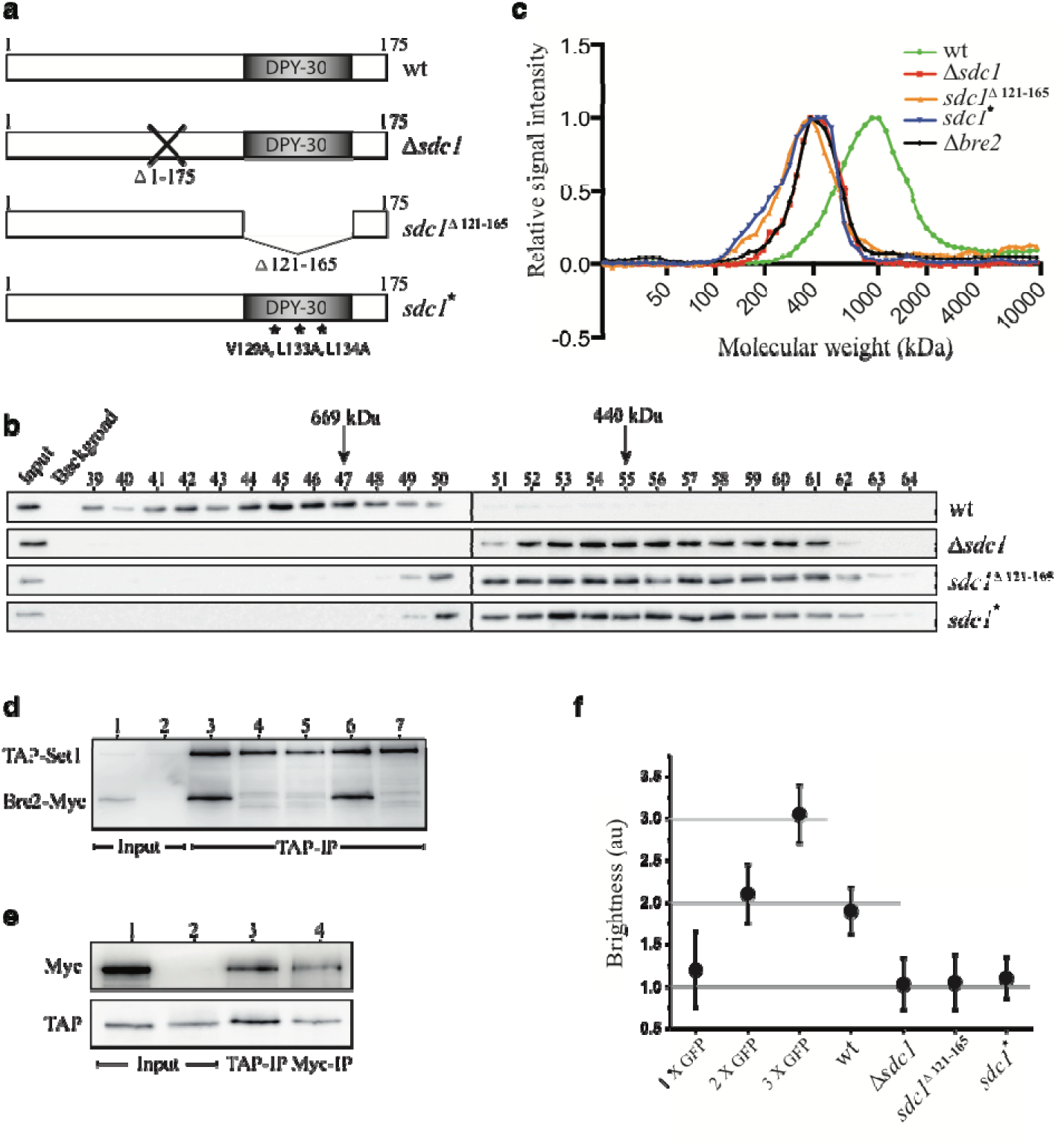
The DPY30 domain of Sdc1 mediates the dimerization of Set1C. (A) Schematic representation of Sdc1 and Sdc1 mutants used in the study showing the conserved DPY-30 box that includes the RIIa dimerization interface, which was deleted (Δ121-165) or point mutated (V129A, L133A, L134A). (B) Size exclusion chromatography on Superose 6 columns of total TAP-Set1 cell extracts from the indicated strains. The columns were calibrated using molecular weight standards and fractions were analyzed by Western for the TAP tag. (C) Plots of the Western analyses from (B) and other size exclusion chromatography runs. (D) Using whole cell extracts from yeast strains expressing TAP-Set1 and Bre2-Myc (lanes 1, 3-6) or just TAP-Set1 (lanes 2, 7), TAP-Set1 was affinity purified with associated proteins (lanes 3-7). The three *sdc1* mutations were present in lane 4 (*Δ*sdc1), lane 5 (*sdc1^Δ121-165^*) and lane 6 (*sdc1\**) and lanes 1 and 2 show input without affinity purification. The Western blot was co-probed with anti-TAP and anti-Myc antibodies. (E) A second copy of the Set1 gene from −497 to +1179 including an N-terminal Myc-tag was integrated into the TAP-Set1 strain at the *ura3* locus (lanes 1, 3, 4). Total cell extract (input) and immunoprecipitates as indicated were analyzed by Western to detect the Myc (top panel) or TAP (bottom panel) tags. Lane 2 – TAP-Set1 only. (F) Photon counting histogram (PCH) analysis of Set1C in whole cell extracts from wt and *sdc1* mutant strains were compared to strains expressing 1x yEGFP, 2x yEGFP (tandem yEGFP-yEGFP) and 3x yEGFP (tandem yEGFP-yEGFP-yEGFP; Supplementary Fig.2). Error bars show mean +/- SEM from three independent experiments.

Because Sdc1 interacts with Bre2 (Roguev et al. 2001; South et al. 2010; Tremblay et al. 2014), we evaluated Set1C lacking Bre2 and found that it was also smaller (Fig. 1C), which is consistent with our previous findings that Set1C lacking Bre2 also lacks Sdc1, and vice versa, but all other subunits remain (Dehé et al. 2006). As expected, complete loss of Sdc1 or deletion of the Dpy-30 box which includes the Bre2 interaction site (South et al. 2010; Tremblay et al. 2014), resulted in concomitant loss of Bre2. However the *sdc1\** triple point mutation, which retains the Bre2 interaction motif, retained Bre2 in Set1C (Fig. 1D). Therefore the Sdc1 dimerization interface and not Bre2 is required for the apparent dimerization of Set1C. These *sdc1* mutations did not alter the expression level of Set1 (Supplemental Fig. 1B) or the association of any other Set1C subunit except Bre2 (Dehé et al. 2006). Notably, because loss of Spp1 alters the H3K4 methylation profile (Morillon et al. 2005; Dehé et al. 2006), we specifically confirmed that Spp1 is retained in the mutant *sdc1* Set1Cs (Supplemental Fig. 1C).

Two further tests for Set1C dimerization were employed. First we made a yeast strain expressing two different tagged versions of Set1: TAP-Set1 and Myc-Set1. Immunoprecipitation using the TAP tag also retrieved Myc-Set1 (Fig. 1E), demonstrating that wild type Set1C includes more than one copy of Set1. Second, we employed fluorescence correlation spectroscopy (FCS) and photon counting for direct quantitation of Set1C stoichiometry in whole cell extracts. Yeast strains were constructed to express yEGFP as a monomer or tandem dimer or trimer from the same locus (Supplemental Fig. 2), which were then used to establish standard values. A yeast strain expressing yEGFP-Swd1 corresponded to the yEGFP dimer whereas the same strain with an *sdc1* mutation corresponded to monomeric yEGFP (Fig. 1F).

**Figure 2.**
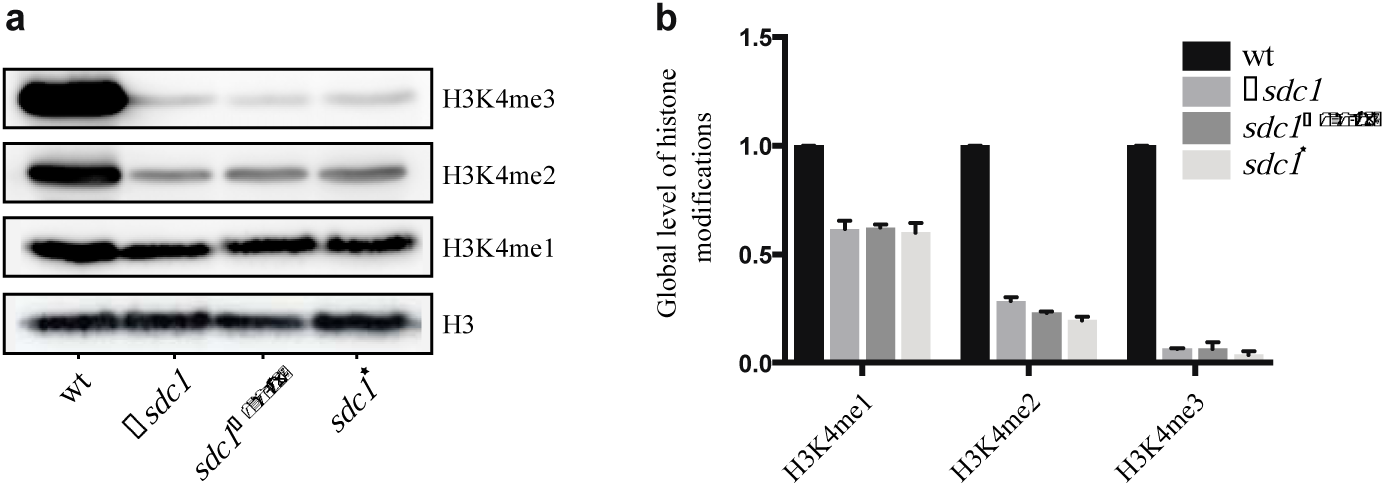
Monomeric Set1C affects global levels of H3K4 methylation. (A) Western blot showing global levels of H3 and H3K4me1, 2 and 3 in wt and *sdc1* mutant strains. (B) Quantification of the signals in (a) and three independent repeats. Error bars show mean +/- SEM from three independent experiments.

Taken together with the 386kDa molecular weight of monomeric Set1C (based on one molecule each of the eight subunits) and the evidence that Sdc1/Dpy30 includes a dimer interface (Wang et al. 2009; Zhang et al. 2015a; Tremblay et al. 2014), these data establish that the approximately 1MDa Set1C is essentially dimeric, Sdc1 presents the dimer interface and the three mutant Sdc1 Set1Cs are monomeric.

### The distribution of H3K4 methylation by monomeric Set1C is skewed

To explore the functional implications of Set1C dimerism, we first compared global H3K4 methylation conveyed by dimeric and monomeric Set1Cs by Western. All three *sdc1* mutations dramatically reduced global trimethylation (H3K4me3), strongly reduced dimethylation (H3K4me2) and moderately reduced monomethylation (H3K4me1) compared to wild type (Fig. 2).

Next, chromatin immunoprecipitation followed by massively parallel DNA sequencing (ChIP-seq) was used to compare the genomic distribution of H3K4me1, 2 and 3 deposited by wildtype dimeric or mutant monomeric Set1Cs. As can be seen in screen shots (Fig. 3A, Supplemental Fig. 3), all three *sdc1* mutations provoked dramatic and virtually identical alterations in the genomic distribution of H3K4 methylation. We first discuss alterations of H3K4me3.

**Figure 3.**
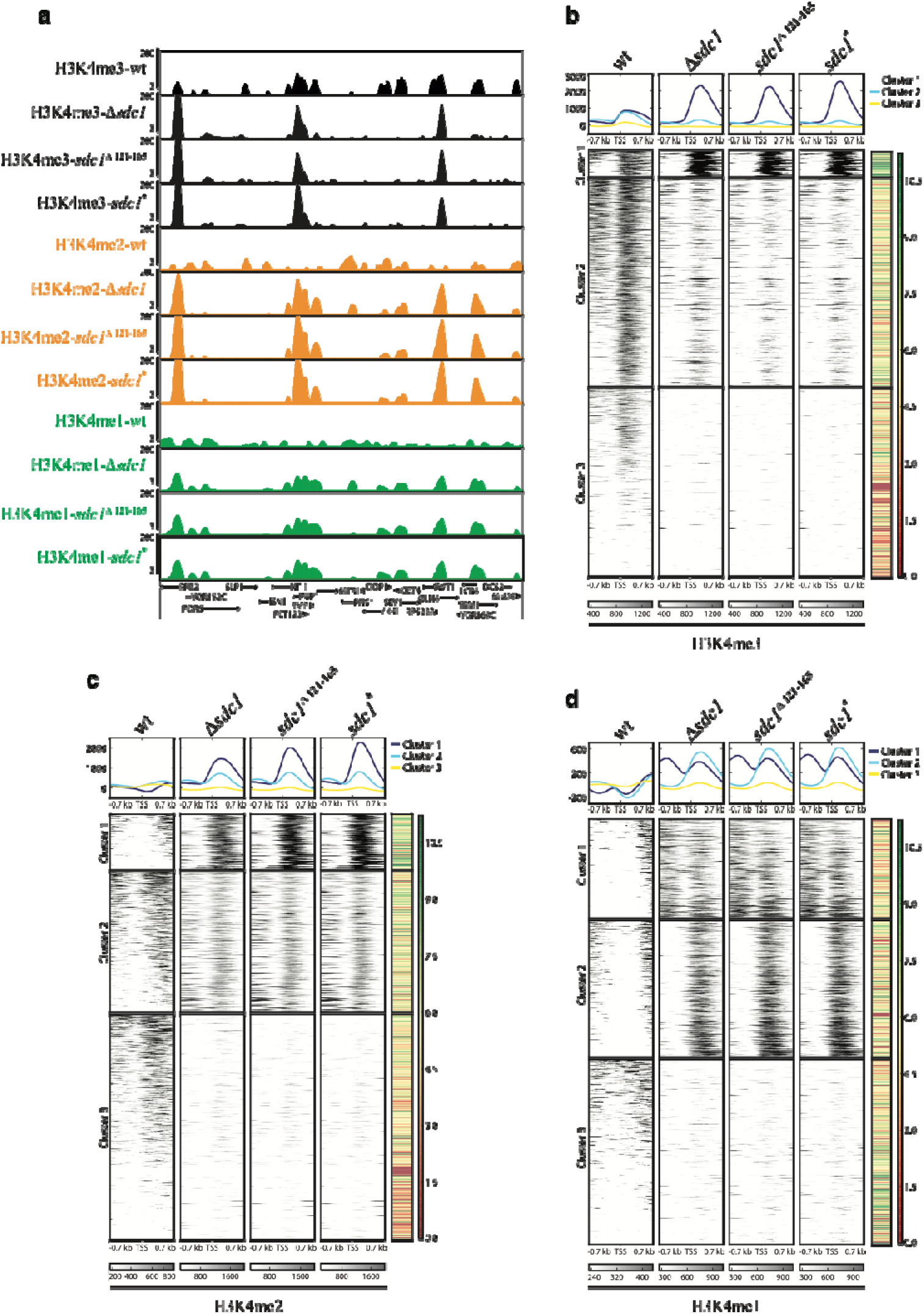
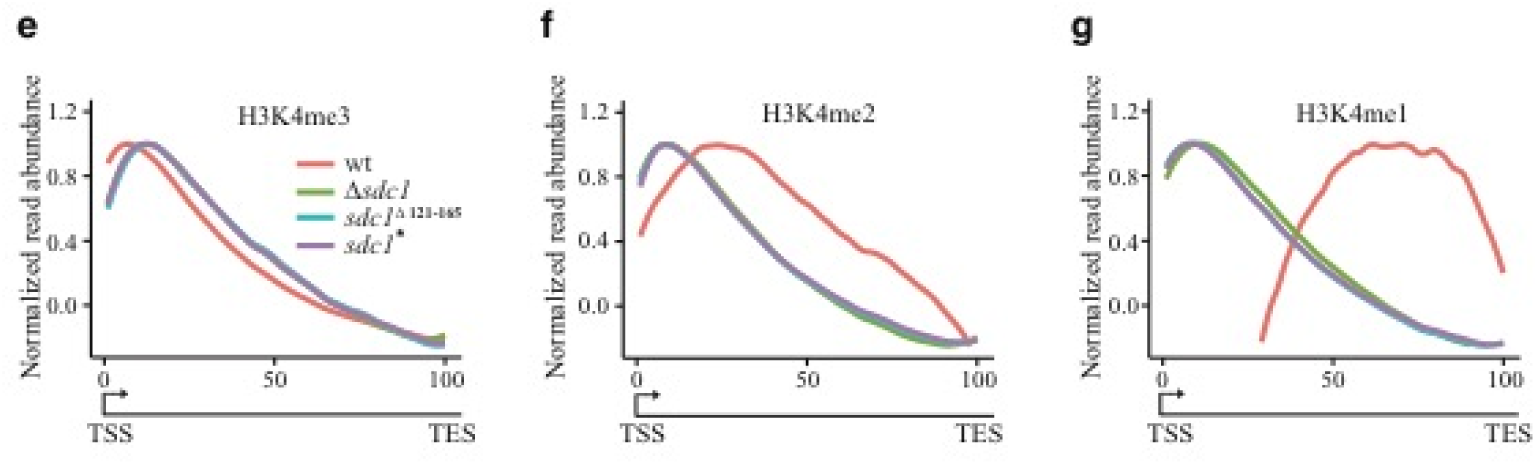
Altered distribution of H3K4 methylation by monomeric Set1C. (A) Screen shot of H3K4me3 (top, black). H3K4me2 (middle, orange) and H3K4me1 (green, bottom) ChIP-seq from wt, *Δ*sdc1, sdc1^Δ121-165^ and *sdc1\** strains as indicated with gene diagrams below. (B) Intensity plots showing H3K4me3 ChIP-seq plotted across a 1.5 kb window centered on transcription start sites (TSSs). K-means clustering generated three clusters (strong, intermediate, weak H3K4me3 intensity) using the *Δ*sdc1 data and then the promoters were stacked according to the strength of the wt H3K4me3 peak downstream of the TSS within each cluster (left hand column; wt). Levels of mRNA expression are shown in the right hand column (colour code at the right; green, high; red, low). At the top of each column, the average profile for each of the three clusters is presented – cluster 1, dark blue; cluster 2, light blue; cluster 3, yellow. (C) As for Fig. 3B except plotting H3K4me2 ChIP-seq. K means clustering and then H3K4me2 stacking according to the peak intensities from *Δ*sdc1 again generated three clusters. These three clusters were independently generated using the H3K4me2 data. Remarkably, the 401 TSSs in cluster 1 of Fig3B (H3K4me3 peaks) are a subset of the 878 TSSs in cluster 1 of Fig 3c (H3K4me2 peaks). (D) As for Fig. 3B except K means clustering of H3K4me1 ChIP-seq with division into three clusters according to bidirectional promoter regions (cluster 1), unidirectional promoter regions (cluster 2) and virtually no H3K4me1 signal (cluster 3). These three clusters do not correspond to the clusters in Fig.3B or C. (E) Metagene profiles of H3K4me3 peaks from wt (red) and *sdc1* mutant yeast strains. The analysis permuted all gene lengths from transcription start site (TSS) to polyadenylation signal (PA) into a scale from 0 to 100. Peak heights were normalized to 1.0. (F) As in (A) except for H3K4me2. (G) As in (A) except for H3K4me1.

Unexpectedly the over ten-fold global reduction in H3K4me3 was not distributed evenly. Only about 7% of all promoters (401 of ~5,500) displayed H3K4me3 peaks and all the rest had almost no H3K4me3 at all. K-means clustering organized the *sdc1* mutations into three clusters with the 401 promoters that retained prominent H3K4me3 peaks conveyed by monomeric Set1C clustered at the top and the promoters totally lacking H3K4me3 at the bottom (Fig. 3B). As repeatedly noted in many eukaryotes, the size of the H3K4me3 promoter peak correlates closely with the amount of mRNA production from that promoter (Howe et al. 2017; Soares et al. 2017). Despite the K means clustering based on the Δsdc1 data, this relationship was approximately retained in the intensity plot of Figure 3B, which shows that the size of the wildtype H3K4me3 peak near the transcription start site (TSS; Fig. 3B; wt column) approximately correlates with mRNA production from most to least (Fig. 3B; far right column).

The distribution of H3K4me2 by monomeric Set1C also related to promoter activity. Normally H3K4me2 is lacking from very active promoters (Soares et al. 2017) (Fig. 3C, cluster 1, wt column). However strongly elevated levels were found at very active promoters in the *sdc1* mutations. Again we used K means clustering to sort the promoters in the mutant *sdc1* strains into three categories according to strongly elevated H3K4me2 (Fig. 3C, top cluster), virtual absence of H3K4me2 (Fig. 3C, bottom cluster) and the rest, which again approximately correlated with mRNA expression levels (Fig. 3C, far right column). Remarkably, the 401 promoters that displayed H3K4me3 peaks (Fig. 3B, cluster 1) are entirely a subset of the 878 promoters showing strongly elevated H3K4me2 (Fig. 3C, cluster 1; Supplemental Fig. 4A).

**Figure 4.**
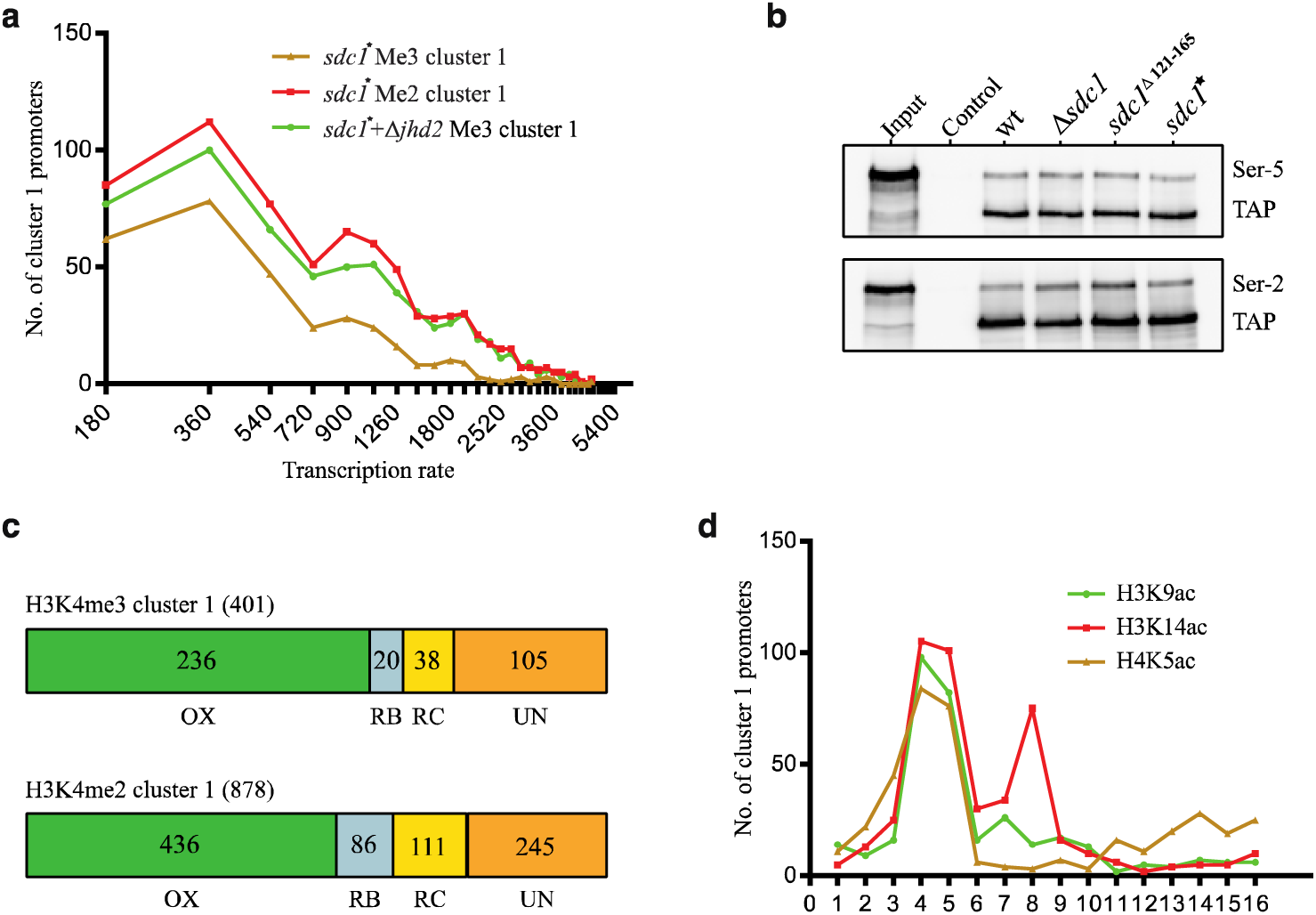
Functional revelations associated with monomeric Set1C (A) The relationship between the 401 promoters that retain H3K4me3 peaks (brown) or the 878 that acquire H3K4me2 peaks (red) in the monomeric Set1C strain was plotted against transcription rates as described by Pelechano et al(Pelechano et al. 2010) in bins of 180 genes from highest to lowest. Also included in this plot is the same analysis for the 787 promoters identified as H3K4me3 rescue peaks upon deletion of Jhd2 (green; see Figure 6). Data is available in Supplementary Table 1. (B) TAP-Set1 was immunoprecipitated from whole cell extracts of wild type and the indicated *sdc1* strains and evaluated by Westerns using antibodies against phosphorylated serine 5 (top panel) or serine 2 (bottom panel) of the Pol II CTD tail together with PAP to visualize the TAP tag, as indicated. Control – IgG sepharose was replaced by sepharose in the immunoprecipitation step. (C) The 401 H3K4me3 and 878 H3K4me2 promoter peaks were assigned to phases of the yeast metabolic cycle (YMC) using the analysis of Kuang et al. (Kuang et al. 2014). OX - oxidative; RB - reductive building; RC – reductive charging phases of YMC. UN – genes unassigned in the YMC data. See also Supplementary Figure 3c,d. (D) The 401 H3K4me3 promoters were compared to the YMC H3K9ac, H3K14ac and H4K5ac ChIP-seq data of Kuang et al. (Kuang et al. 2014). The plot shows the number of 401 promoters that occur in the top 500 promoters displaying the nominated acetylation in each of the 16 ChIP-seq samples that were taken across the YMC. Samples 3, 4 and 5 are in the oxidative phase.

The distribution of H3K4me1 was also distorted by the *sdc1* mutations (Fig. 3D), however the relationship to mRNA expression level was less obvious. Normally H3K4me1 is lacking from all active promoters however more than half of all promoters in the *sdc1* mutant strains displayed some H3K4me1, which we divided into two categories according to double (top cluster, bidirectional promoters) or single (middle cluster) peaks.

Notably, no differences in H3K4 methylation conveyed by monomeric Set1C with or without Bre2 (i.e. *Δ*sdc1 or *sdc1^Δ121-165^* compared to point mutated *sdc1\**) were observed. This may indicate that Bre2 is not required for H3K4 methylation. However Bre2 may still contribute *in vivo* in the absence of Sdc1 through a loose association that was lost upon our biochemical fractionation. Relevant to this possibility, the human homologue of Bre2, ASH2L, not only associates with DPY30 but also with the human homologue of Swd1, RbBP5 (Avdic et al. 2011; Zhang et al. 2015b).

### Shifted H3K4me2 and 1 deposition by monomeric Set1C

The H3K4me ChIP-seq data were used to generate metagene profiles (Fig. 3E-G). For the 401 genes that displayed H3K4me3 peaks, the H3K4me3 distribution was essentially the same as wild type with the peak on the first transcribed nucleosome and diminishing distribution along the transcribed regions. However global H3K4me1 and 2 profiles were strongly shifted towards the TSS suggesting that the absence of H3K4me3 allowed H3K4me1 and 2 to occupy normally excluded positions.

### Correlation to transcription rate and the oxidative phase of the yeast metabolic cycle

To examine the relationship between the 401 H3K4me3 cluster 1 genes and transcription more closely, we used the mRNA expression levels and transcription rate estimates made by Pelechano et al (Pelechano et al. 2010) to find that two-thirds (265/401; 66%) are amongst the 1000 most highly expressed genes (Supplemental Table 1). The relationship between the most highly expressed genes and the 401 H3K4me3 cluster 1 genes was plotted using bins of 180 genes from highest to lowest transcription rate (Fig. 4A). The same analysis for the 878 H3K4me2 cluster 1 genes also displayed a strong relationship to the transcription rate (Fig. 4A).

In wild type yeast, Set1C associates with elongating RNA Polymerase II, so we evaluated whether monomeric Set1C interaction with Pol II had been altered. However no change was observed. As previously reported for wt Set1C (Dehé et al. 2006), immunoprecipitation of TAP-tagged wt and monomeric Set1 retrieved both serine 5 and serine 2 phosphorylated forms of Pol II (Fig. 4B).

GO term analysis of the 401 genes revealed highly statistically significant relationships especially relating to ribosome biogenesis (Supplemental Fig. 4B), which is not surprising because the genes involved in ribosome biogenesis are amongst the most highly expressed in yeast. Ribosome biogenesis is also the major feature of the oxidative phase of the yeast metabolic cycle (YMC; {Tu:2005cv}). Therefore we checked the relationship of the 401 genes to a high resolution analysis of the YMC (Kuang et al. 2014). A strong correlation to genes in the oxidative phase (OX) emerged (Fig. 4C, Supplemental Fig. 4C) despite the fact that OX is the shortest phase occupying 1/5^th^ of the YMC with the other two phases, reductive building (RB) and reductive charging (RC), together occupying 4/5^th^. Nearly 60% of the 401 genes (236/401) are OX genes with 72/236 distinct from the 265 in the top 1000 highly transcribed genes. Furthermore, a strong correlation between the 401 genes and promoters displaying the three histone acetylation marks that peak in the oxidative phase (H3K9ac, H3K14ac, H4K5ac (Kuang et al. 2014) was also observed (Fig. 4D). The relationship of the 878 H3K4me2 cluster 1 genes to the oxidative phase is also strong (Fig. 4C, Supplemental Fig. 4E).

### H3K4me3 peaks deposited by monomeric Set1C are reduced compared to wt

In the intensity plots and screen shots of Figure 3, all H3K4me ChIP peaks in the *sdc1* mutant strains appear to be stronger than their wild type counterparts. However the immunoprecipation efficiencies from wt and mutant chromatin, using the same antibody and genome inputs, are not comparable because the ratios of antibody to epitope are very different. To address this variance and determine a quantitative adjustment factor, we repeated the ChIPs with an input control using wild type *S. pombe* chromatin. Adjusting to the input control indicated that the H3K4me3 peaks in the *sdc1* mutants should be reduced about eight-fold (Supplemental Fig. 5), thereby indicating that the H3K4me3 peaks in the monomeric Set1C strains were actually strongly reduced compared to their wild type counterparts. Similarly, H3K4me2 peaks should be reduced about three-fold. However the adjustment factors for H3K4me1 indicated that yields should be slightly increased. These adjustment factors concur with expectations based on the global Western analysis (Fig. 2).

**Figure 5.**
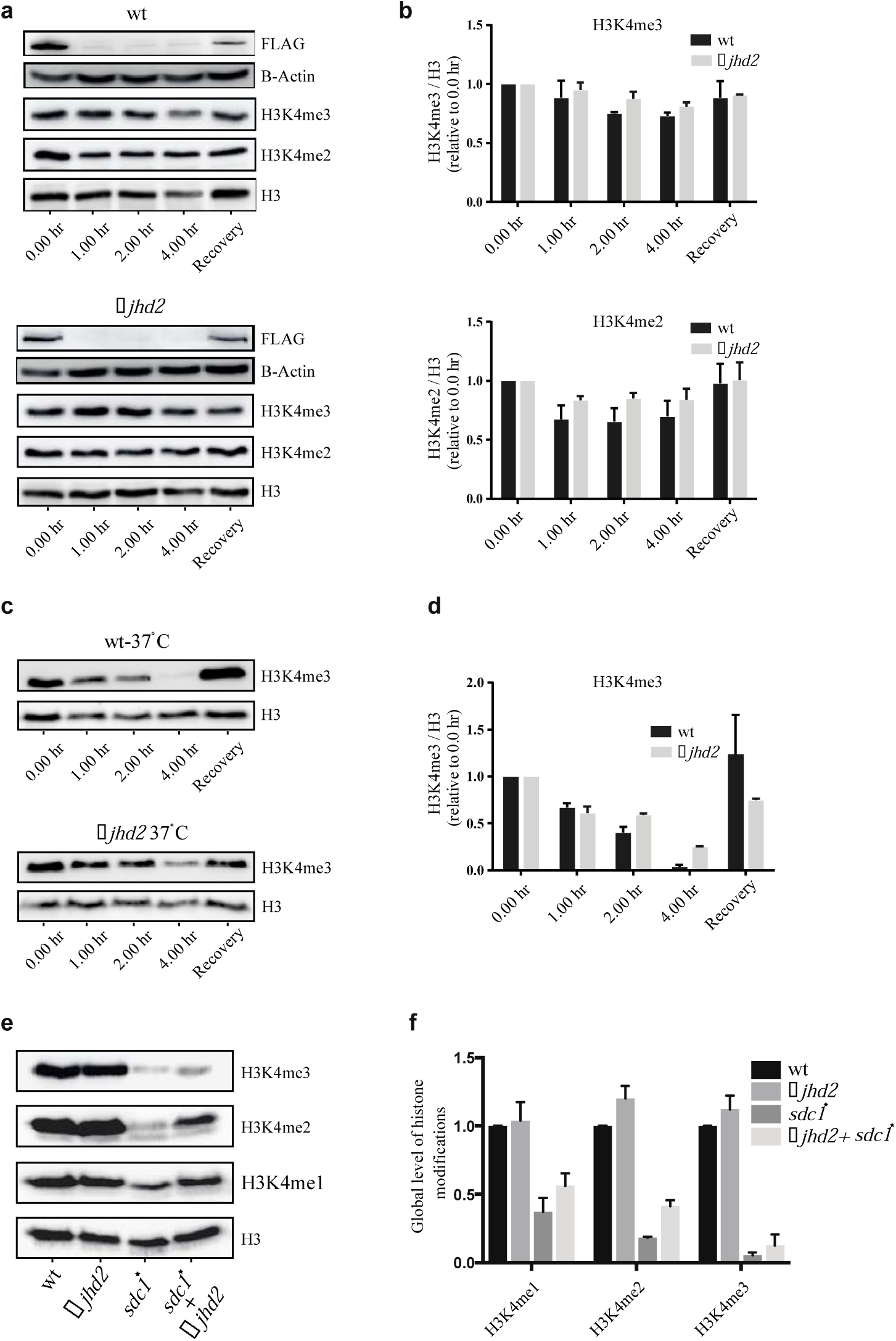
Demethylation by Jhd2 (A) Time courses after IAA administration in a strain expressing OsTir1 with AID-Set1 (upper panel) and AID-Set1/*Δ*jhd2 (lower panel). Western analyses were performed with antibodies against the epitopes indicated at the right. Recovery was harvested 2 hours after IAA washout. (B) Quantification of H3K4me3 (upper histogram) and H3K4me2 (lower histogram) of the experiment in (A) and two repeats, normalized to H3 levels. Error bars; mean +/- SEM from 3 independent experiments. (C) The same experiment as (B) performed at 37°C. (D) Quantification of H3K4me3 from the experiment in (C) as described in (B). (E) Western blot showing global levels of H3 and H3K4me1, 2 and 3 in wt, *Δ*jhd2, (F) *sdc1\** and *sdc1*+Δjhd2* strains. (F) Quantification of the signals in (E) and two repeats. Error bars, mean +/- SEM.

### Alterations of gene expression and H3K4 methylation do not correlate

Total mRNA profiles were documented to evaluate the impact of monomerizing Set1C on gene expression. Because complete removal of H3K4 methylation has only a modest effect on gene expression in yeast (Margaritis et al. 2012; Howe et al. 2017; Ramakrishnan et al. 2016), we were not surprised to find that these dramatic alterations in the distribution and levels of H3K4 methylation also had modest effects on gene expression (Supplemental Fig. 6). Overall only about 4% (211) of all mRNAs were commonly affected more than two fold in all three *sdc1* mutants (Supplemental Fig. 6C,D) and these alterations were evenly spread across expression levels and all three H3K4me3 clusters (Supplemental Fig. 6G-I), so did not correlate with promoter activity. Notably, the 134 commonly affected genes whose expression increased in the three *sdc1* mutant strains were enriched for ribosomal biogenesis (Supplemental Fig. 6F), which is reminiscent of the apparent role of Set1 as a repressor of ribosomal genes (Ramakrishnan et al. 2016). Most of these genes are not in the 401 genes of H3K4me3 cluster 1 (105/134; 78%) or the 878 genes of H3K4me2 cluster 1 (79/134; 59%) again indicating that alterations of H3K4 methylation and expression did not correlate.

**Figure 6.**
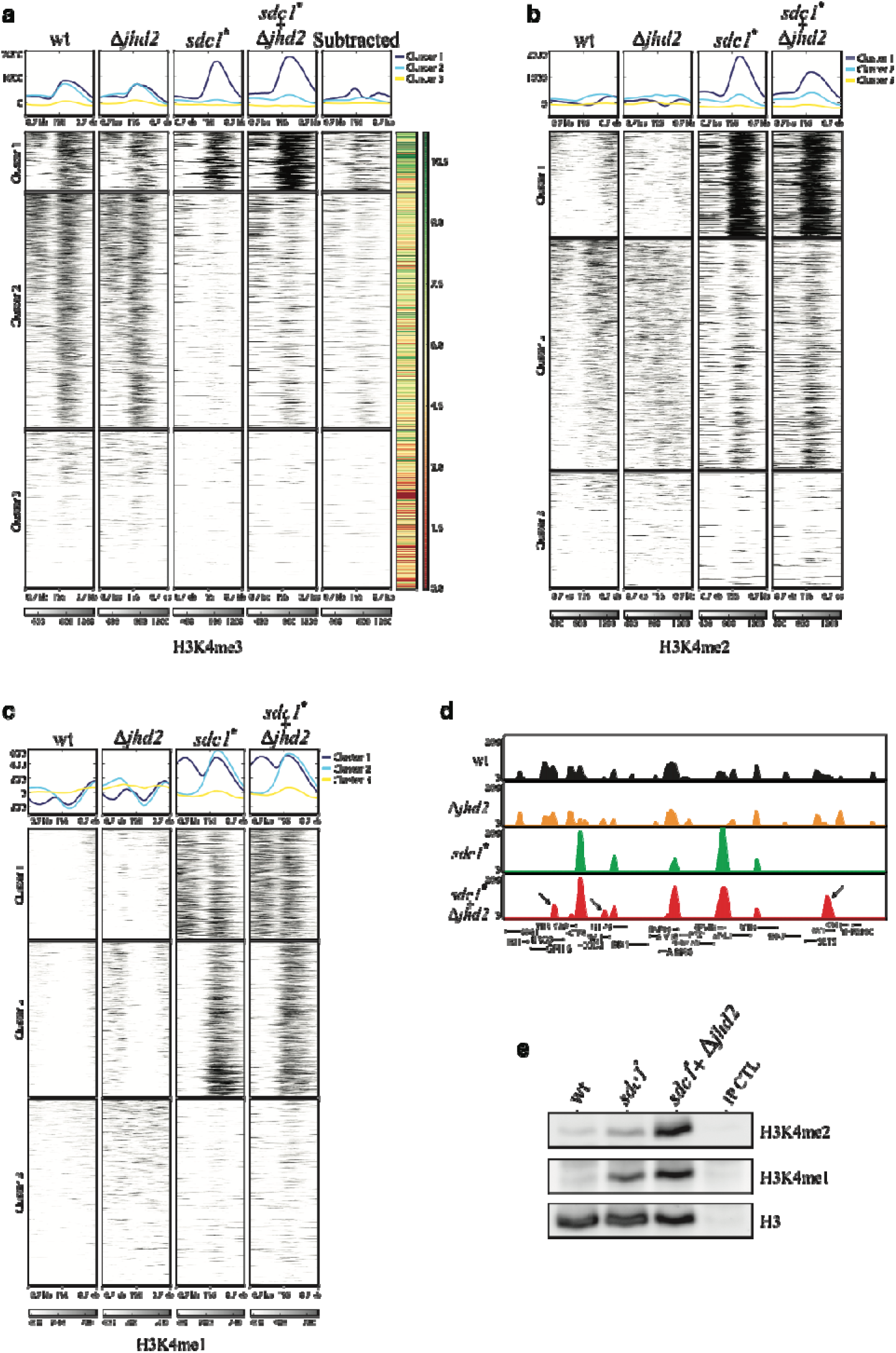
Jhd2 demethylation of asymmetric H3K4me3 deposited by monomeric Set1C (A) As for Figure 3B, intensity plots display ChIP-Seq enrichment for H3K4me3 plotted across a 1.5 kb window centered on transcription start sites (TSS). The three clusters identified by K means are the same as Figure 3B. The ‘Subtracted’ column is the result of subtracting column ‘*sdc1*’* from column ‘*sdc1* + Δjhd2’* to illustrate the effect of Jhd2 demethylation. mRNA expression levels are shown in the right hand column (colour code at the right). (B) As for Figure 6A, intensity plots showing ChIP-Seq enrichment for H3K4me2 plotted across a 1.5 Kb window centered on transcription start sites (TSS). The three clusters identified by K means are the same as Figure 3c. (C) As for Figure 6A, intensity plots showing ChIP-Seq enrichment for H3K4me1 plotted across a 1.5 Kb window centered on transcription start sites (TSS). The three clusters are the same as Figure 3D. (D) Screen shot of H3K4me3 profiles from the indicated strains. Arrows indicate some peaks that appear when *jhd2* was mutated in the *sdc1\** strain. (E) H3K4me3 mononucleosomes were immunoprecipitated from wt, *sdc1\** and *sdc1*+Δjhd2* strains and analyzed for H3K4me1 and 2 normalized to H3 by Western. The H3K4me3 antibody was omitted from the immunoprecipitation control (IP CTL). Full gel images are available in S8C.

### Demethylation by Jhd2 in normal growth is modest

The altered patterns of H3K4 methylation in the monomeric Set1C strains could also be affected by demethylation mediated by the sole yeast jumonji-domain H3K4 demethylase, Jhd2 (Liang et al. 2007; Huang et al. 2010). Before we could address this issue, we first needed to address the controversy surrounding the demethylation activity of Jhd2 in wild type yeast. Initially loss of H3K4me3 was reported to be passive through dilution by replication (Ng et al. 2003). Subsequently, a much higher rate of quantitative, Jhd2-dependent, H3K4me3 removal within one cell cycle was reported using a Set1 temperature-sensitive degron (Seward et al. 2007). A more detailed analysis identified both active and passive contributions to demethylation (Radman-Livaja et al. 2010).

To determine whether Jhd2 rapidly demethylates H3K4me3 under normal laboratory growth, we employed the auxin inducible degron (AID; (Nishimura et al. 2009) by adding it onto the N-terminus of Set1. As performed with the temperature-sensitive degron (Seward et al. 2007), yeast cultures were treated with α-factor to synchronize and reduce cycling before degron activation by indole acetic acid (IAA) administration. AID-Set1 was rapidly degraded upon IAA addition accompanied by modest reductions of H3K4me2 and 3 levels, which were partly attributable to Jhd2 (Fig. 5A,B). Notably, the rate of H3K4me3 loss after auxin degron removal of Set1 was much less than that reported using a temperature-sensitive degron (Seward et al. 2007). Because the activity of chromatin regulators is often amplified by stress (Weiner et al. 2012), we considered the possibility that the heat shock employed for the Set1 temperature-sensitive degron was the source of the discrepancy. Hence the auxin degron experiment was repeated to include a shift to 37°C at the time of IAA administration, which revealed rapid H3K4me3 demethylation, some of which was mediated by Jhd2 (Fig. 5C,D). Because temperature shift to 37°C releases yeast from α-factor blockade (Day et al. 2004), passive dilution through histone replacement by cycling could have also contributed to the loss of H3K4me3 in this experiment, as well as other potential mechanisms such as tail cleavage (Duncan et al. 2008; Santos-Rosa et al. 2009). We conclude that H3K4me3 and 2 demethylation is modest under normal culture conditions and is promoted by heat shock through Jhd2 and another active mechanism.

### Jhd2 removes H3K4me3 from many sites of monomeric Set1C action

Consistent with this conclusion and previous studies, deletion of *jhd2* in wild type yeast under normal growth conditions had only slight effects on total H3K4 methylation (Fig. 5E.F), gene expression (Supplemental Fig. 6A,B; 7) or the distribution of H3K4 methylation (Fig. 6 *Δ*jhd2 columns; Supplemental Fig. 8). In contrast to these subtle effects, loss of Jhd2 had a pronounced impact on H3K4me3 in the monomeric Set1C yeast strain, which was elevated in both clusters 1 and 2 to increase peak sizes and reveal many (787) previously absent peaks (Fig. 6A,D). This reveals that a substantial amount of H3K4me3 deposited by monomeric Set1C is removed by Jhd2. In contrast, loss of Jhd2 resulted in only slight alterations of the H3K4me1 and 2 patterns (Fig. 6B,C).

The 787 “rescued” H3K4me3 peaks strongly overlap with the previously defined categories with 87% (684) also H3K4me2 cluster 1 genes (Supplemental Fig. 4A). The rescued peaks clearly also relate to the transcription rate (Fig. 4A) and the YMC oxidative phase (Supplemental Fig. 4F).

Transcriptome profiling evaluated the impact of *jhd2* deletion in combination with the *sdc1* point mutation. Deletion of *jhd2* had no effect on wild type gene expression and a modest impact when combined with point mutated *sdc1* (Supplemental Fig. 7). Despite robust data quality (Supplemental Fig. 6A,B), of the 558 mRNAs that were altered more than 2 fold in the point mutated *sdc1* strain, 327 (59%) were similarly altered in the double mutant strain (Supplemental Fig. 7A,B). Again, these gene expression changes were evenly distributed regardless of expression levels or promoter activity from strong to weak (Supplemental Fig. 7E-G) and consequently did not correlate with H3K4me3 alterations.

**Figure 7.**
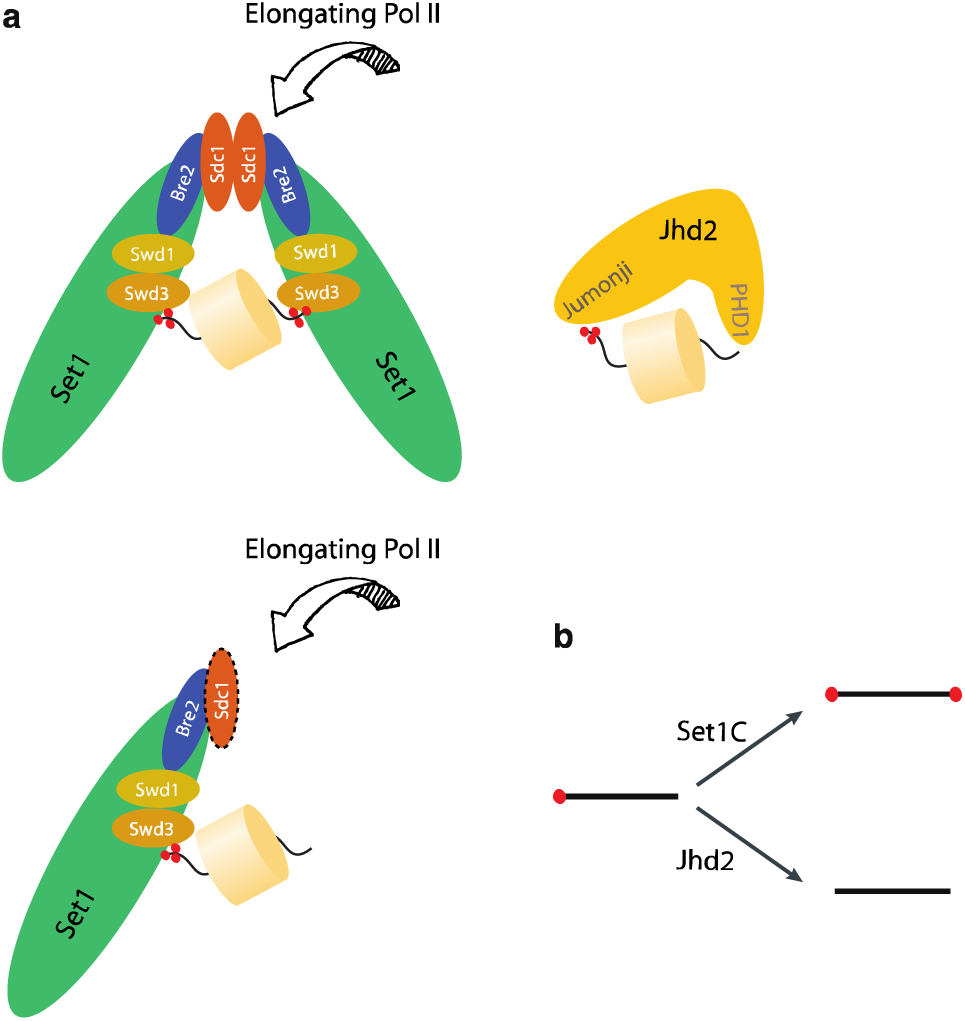
Model for action by Set1C and Jhd2. (A) Wild type Set1C is dimeric and is introduced by elongating Pol II to symmetrically trimethylate promoter nucleosomes at H3K4. Other factors and cues modulate this process. The Sdc1 dimer interface associates with Set1C through interaction with Bre2. The other WRAD heteromer, Swd1/Swd3 is required for methylation (Wdr5 = Swd1; Rbbp5 = Swd3; Ash2l = Bre2; Dpy30 = Sdc1). When Sdc1 is mutated, Set1C is monomeric. It is still introduced by elongating Pol II to promoter nucleosomes to trimethylate H3K4 but only on one tail leaving asymmetrically modified nucleosomes. Asymmetric H3K4me3 nucleosomes are de-trimethylated by Jhd2 through recognition of the un/mono-methylated tail by its’ PHD1 finger. (B) Dimeric Set1C and monomeric Jhd2 act in concert to reduce asymmetrical H3K4me3.

### Monomeric Set1C produces increased levels of asymmetric H3K4me nucleosomes

As recently evaluated (Soares et al., 2017) and also observable in Figure 3B-D, H3K4me2 and 1 are excluded from the H3K4me3 nucleosomes that surround promoters in wild type yeast and the degree of exclusion relates to the transcriptional activity of the promoter. For highly expressed genes, both H3K4me2 and 1 are completely excluded. For moderately expressed genes, H3K4me1 but not H3K4me2 is excluded. For lowly expressed genes, both H3K4me1 and 2 infiltrate promoter nucleosomes (Fig. 3B-D, wt panels). Monomeric Set1C changed the patterns with H3K4me3 reduced on all promoters whereas H3K4me1 and 2 are increased on active promoters (Fig. 3B-G; 6A-C; Supplemental Fig. 8). To evaluate whether this represents a shift from symmetrical H3K4me3 nucleosomes to asymmetrical H3K4me3 combined with H3K4me1 or 2, we scaled up the ChIP procedure four-fold using mononucleosomes (Supplemental Fig. 9) and analyzed the H3K4me3 immunoprecipitates by Western for H3K4me1 and 2. This analysis clearly revealed elevated levels of asymmetrically H3K4me modified nucleosomes in the monomeric Set1C strains (Fig. 6E).

## DISCUSSION

Until recently, the inherent symmetry of the nucleosome promoted the assumption that post-translational modifications would also be symmetrical. The identification of asymmetrically modified nucleosomes at active promoters (Rhee et al. 2014; Voigt et al. 2012) revealed a new level of epigenetic detailing, which could convey greater combinatorial specificities and/or impose directional orientation in chromatin. One hit stochastic deposition was the first proposition for the origin of nucleosomal post-translational asymmetry (van Rossum et al. 2012). Our finding that Set1C is dimeric introduces the new perception that deposition of H3K4 methylation is inherently symmetrical (Fig. 7). This proposition has several implications that we first discuss in the light of the recent high resolution structures of the Set1C enzymatic core (Hsu et al. 2018; Qu et al. 2018).

Both the recent crystal and cryo-EM structures present a monomeric Set1C/COMPASS complex centered on the C-terminal SET domain of Set1 co-purified with the conserved WRAD quartet of Swd3, Swd1, Bre2 and Sdc1. In both structures, all five components are monomeric except Sdc1, which is a dimer based on the RIIa dimer interface. One Sdc1 interacts with Bre2 though the Bre2 interaction motif and the other Bre2 interaction motif is exposed (Hsu et al. 2018). Consequently dimeric Set1C can be readily accommodated with these high resolution structures, possibly with the two monomeric complexes forming a cup that can encompass a nucleosome rather than pointing away from each other in opposite directions. The monomeric high resolution structures concord with the genetic/mass spectrometry dissections of the complex (Roguev et al. 2001; Dehé et al. 2006) including the positioning of Spp1 (Qu et al., 2018) with one notable exception. Removal of Sdc1 also results in loss of Bre2 from Set1C. The high resolution monomeric structures do not offer an explanation for this finding. We suggest that the explanation lies with the role of the Bre2-Sdc1 heteromer (Roguev et al. 2001) in Set1C dimerization. From the differences between their two cryo-EM structures, Qu et al (2018) speculate that the flexible curvature of their complex could be binding site to accommodate a nucleosome. We extend this idea to suggest that this flexible curvature binds a nucleosome on one side and this binding is reciprocated by dimeric Set1C to bind both sides. Consequently, the study of Set1C nucleosomal binding needs to be performed with the dimeric complex.

Mutagenesis of the Set1C dimer interface enabled a ChIP-seq comparison of the genomic H3K4 methylation distribution in wild type dimer and mutant monomeric Set1C yeast strains. Importantly, mutagenesis of the dimer interface did not interfere with the association of Set1C with elongating RNA Pol II. Based on this association and the widespread correlation between mRNA levels and the size of H3K4me3 promoter peaks, it is widely presumed that H3K4me3 deposition on promoter nucleosomes follows transcription of the associated gene (Howe et al. 2017; Woo et al. 2017). Whereas H3K4me3 peaks in wild type yeast smoothly relate to the amount of mRNA produced, only 401 promoters displayed H3K4me3 peaks in the monomeric Set1C strains. Furthermore in the monomeric Set1C yeast strains, both H3K4me2 and 1 infiltrated active promoters from which they are normally excluded. The 401 H3K4me3 promoters are a subset of the 878 promoters that showed strong H3K4me2. Both the 401 and 878 genes are strongly related to high transcription rates and the YMC oxidative phase.

Removal of the sole yeast H3K4 demethylase, Jhd2, provided insight. Whereas Jhd2 removal in a wild type background had virtually no effect on H3K4 methylation patterns or gene expression, Jhd2 removal from a monomeric Set1C strain resulted in increased H3K4me3 both on the 401 promoters and also on another 787 promoters, which were largely (685/787=87% or 685/878=78%) the same as the 878 H3K4me2 promoters. Therefore certain mechanisms are directing the monomeric Set1C to a subset of promoters resulting in a core that display H3K4me3 and a broader category that are demethylated by Jhd2 and display H3K4me2. These mechanisms correlate with high transcriptional activity and the YMC oxidative phase, possibly through recognition of acetylations at H3K9, H3K14 and/or H4K5. However neither of these correlations appear to be comprehensive. Potentially another factor or combination of factors is required to fully account for the promoter selectivity of monomeric Set1C.

Normally, H3K4me3 deposited by dimeric Set1C is not removed by Jhd2 whereas a substantial fraction of H3K4me3 deposited by monomeric Set1C was removed by Jhd2. Therefore H3K4me3 deposited by monomeric and dimeric Set1Cs differ in some respect. Immunoprecipitation of H3K4me3 mononucleosomes indicated that increased H3K4me asymmetry is the notable difference.

Jhd2 is expressed as a monomer without associated proteins (Liang et al. 2007). Furthermore recent *in vitro* evidence using KDM5A, one of the human homologues of Jhd2, revealed that PHD1 zinc finger, which is highly conserved in Kdm5 demethylases including Jhd2, binds to the H3 tail with the preference H3K4me0>me1>me2>me3 and when bound stimulates H3K4me3 demethylation by the Jhd2 Jumonji domain (Torres et al. 2015). Evidence from KDM5B also supports a key role for PHD1 in binding H3K4me0 and demethylation (Zhang et al. 2014; Klein et al. 2014). We extend these observations to suggest that stimulation is exerted when PHD1 binds one H3 tail on asymmetrical H3K4me3 nucleosomes (Fig. 7A).

The combination of symmetrical H3K4 trimethylation conveyed by dimeric Set1C with asymmetric H3K4me3 demethylation conveyed by monomeric Jhd2 provokes a remarkable conclusion. Rather than methylation and demethylation acting in opposition as logic would suggest, the two processes act in concert to reduce asymmetric H3K4me3 nucleosomes and thereby focus symmetry onto selected nucleosomes (Fig. 7B). A juxtaposition of concerted addition and singleton removal could also apply to create symmetry and organization on other symmetrical substrates and circumstances.

This concept also provides some clarity regarding the action of Kdm5/Jarid 1 class of demethylases, which may have relevance for other classes of histone demethylases. As reported by others (Ramakrishnan et al. 2016; Tu et al. 2007) and again documented here, loss of Jhd2 has almost no effect in yeast grown under normal laboratory conditions. Similarly, mouse knock-out phenotypes of the Kdm5/Jarid1 genes have been difficult to interpret because they present incompletely penetrant phenotypes (Schmitz et al. 2011; Scandaglia et al. 2017). Concordant with studies on Kdm5b in embryonic stem cells (Kidder et al. 2014), we suggest that an ancillary editing role in the epigenetic definition of chromatin offers an explanation for the incompletely penetrant phenotypes of these highly conserved enzymes. Under optimal circumstances their contribution is unnecessary but in certain circumstances the demethylases serve to reduce errors. That is, the Kdm5/Jarid1 demethylases add correctional stability to gene expression programs by focusing H3K4me3 onto promoter nucleosomes.

Similarly, the role of Set1 and H3K4 methylation in yeast has been a source of confusion largely due to its apparent dispensability and the counter intuitive observation that more genes show increased rather than decreased expression when Set1 is removed (Margaritis et al. 2012; Lenstra et al. 2011). Our observations add more confidence to the presumption that H3K4me3 on promoter nucleosomes is a consequence of the association of Set1C with the elongating polymerase (Howe et al. 2017; Woo et al. 2017; Soares et al. 2017). Given the conserved role of the H3K4me3 epitope as a binding site for protein complexes involved in transcriptional initiation such as TFIID (Vermeulen et al. 2007), a contribution to the stability of gene expression, rather than a role in its regulation, appears to be the most robust explanation.

Along with TFIID and other complexes involved in transcriptional initiation, H3K4me3 is also bound by Set1C itself through the PHD finger in Spp1 (Shi et al. 2007; Eberl et al. 2013). Spp1 binding of H3K4me3 could serve as a feed-forward epigenetic mechanism to propagate H3K4me3 after replication. Alternatively, our new perceptions about H3K4me3 symmetry on promoter nucleosomes suggest that Spp1 could serve in other ways. If delivery of Set1C to the promoter nucleosomes by its association with elongating polymerase resulted in one-hit, stochastic, deposition then Spp1 binding could serve to attach Set1C to the hemi-trimethylated nucleosome for trimethylation of the other tail. Another possibility involves an allosteric role for Spp1 binding within the dimeric complex to enhance the efficiency of symmetrical trimethylation.

We favour the second explanation because the difference between H3K4me3 deposition by monomeric or dimeric Set1Cs suggests that monomeric Set1C deposition is stochastic whereas dimeric Set1C deposition includes co-operativity to which Spp1 binding to H3K4me3 may contribute. This possibility concords with the cryo-EM structure, which includes Spp1. Although the PHD finger was not resolved, the authors suggest that it’s proximity may permit a contribution to substrate binding (Qu et al. 2018).

We suggest that in response to transcriptional activity and subsequent verification according to oxidative phase and other unidentified cues, dimeric Set1C symmetrically trimethylates two H3K4 residues on the same tail of promoter nucleosomes. Deposition by monomeric Set1C also reflects transcriptional activity and the other verification cues however it does not engage in dual deposition but rather one-hit action resulting in asymmetrically trimethylated nucleosomes. When the transcription rate and/or other cues are higher than a certain level, sufficient stochastic deposition by monomeric Set1C achieves symmetrical trimethylation of promoter nucleosomes, which then resist demethylation by Jhd2 because it is symmetrical.

We also suggest that the dimeric Set1C, which is about four times larger than a nucleosome, may have a specific nucleosomal binding pocket that serves to present the two histone 3 tails to the SET domains. Our suggestion concords with nucleosome binding speculations promoted by the monomeric cryo-EM structure, which we extend by demonstrating that Set1C is dimeric. Our observations do not permit conclusions about symmetry regarding H3K4me1 and 2 deposition. However, if dimeric Set1C binds to a nucleosome and accesses both H3 tails, catalytic symmetry for mono- and di-methylation is also likely.

The SET domain-WRAD scaffold is amongst the most highly conserved protein modules in eukaryotic epigenetics and all H3K4 methyltransferase complexes appear to include Dpy30/Sdc1 (Ruthenburg et al. 2007; Rao and Dou 2015). Therefore it is likely that all Set1/Trithorax-type H3K4 methyltransferase complexes are dimeric and their deposition of methyl groups in chromatin is, like Set1C, also an embedded product of dimerism. This implies that H3K4me3 nucleosomes are implicitly symmetrical for H3K4me3 and asymmetrical H3K4me3 nucleosomes, such as bivalent nucleosomes (Voigt et al. 2012; 2013; Shema et al. 2016), are the product of asymmetric demethylation or interference with the propensity of the bivalent H3K4 methyltransferase, Mll2 (Denissov et al. 2014), to methylate both H3 tails in the same nucleosome.

Using the auxin degron, we observed that bulk H3K4me3 and 2 turnover is modest. Attempting to understand previous results (Seward et al. 2007), we found that H3K4me3 loss is promoted by heat shock, which may contribute to the accompanying widespread changes in gene expression. Notably, less than half of the H3K4me3 loss could be attributed to Jhd2 and another mechanism for demethylation is clearly indicated. Under stress such as heat shock, it appears that symmetrical H3K4me3 nucleosomes can be disturbed possibly to generate asymmetrical H3K4me3 nucleosomes that Jhd2 acts upon. Alternatively, heat shock may activate Jhd2. In either case, examination of the impact of stress on demethylation could be fruitful.

## Materials and Methods

Additional methods are described in detail in the Supplement.

### Yeast Strains

Strains used in this study are listed in Supplementary Table 1. Set1 was N-terminally tagged with an AID-FLAG cassette because C-terminal tagging inactivates the methyltransferase activity (Roguev et al. 2001). The AID-FLAG cassette was directed between the 2^nd^ and 3^rd^ amino acid codons of the Set1 gene by insertion of KanMx selection gene flanked by loxP sites and selection for G418 resistance. After Cre recombination, a 34 bp loxP site was left in the reading frame between AID-FLAG and the rest of the Set1 coding region.

### Protein assays and immunoblotting

For size exclusion chromatography, a 10/30 Superose 6 size exclusion column (HR, Pharmacia) was loaded with 500μl of cleared crude cell extract from a TAP-tagged strain and run in glycerol-free buffer E (20 mM HEPES-NaOH, pH 8.0, 350 mM NaCl, 0.1% Tween 20, 10% glycerol, 1ug each of leupeptin, aprotinin, and pepstatin A, 1 mM phenylmethylsulfonyl fluoride (Logie and Peterson 1999). Fractions were resolved on an 8% SDS-PAGE gel and analyzed by immunoblotting with peroxidase-anti-peroxidase (PAP, Sigma) diluted 1:1000, for detection of the protein A region within the TAP tag using the ECL kit (Amersham Pharmacia Biotech).

### Antibodies

α-Myc antibody (Myc; 9E10, catalog number 11667203001, Roche Applied Science); PAP antibody (catalog number P1291, Sigma); H3K4me1 (pAB- 037-050, 1:2000, Diagenode); H3K4me2 (pAB-035-050, 1:1000, Diagenode); H3K4me3 (ab8580, 1:1000, Abcam); H3 (ab1791, 1:2000, Abcam); CTD Ser2 phosphorylation (Abcam, ab5095; 1:1000); CTD Ser5 phosphorylation (Abcam, ab5131; 1:1000).

### Fluorescence Correlation Spectroscopy (FCS)

Swd1 was C-terminally tagged with yeast codon optimized EGFP. To generate yeast strains expressing yEGFP as a monomer or tandem dimer or trimer, the coding sequence of yEGFP was PCR amplified with a forward primer containing the initiating codon along with a 5’ 40bp homology arm to the integration vector, YIplac211, and a reverse primer containing HindIII and BamHI sites along with a 3’ 40bp homology arm. Linearized YIplac211 and the PCR fragment were then recombined by full length RecE/RecT recombination (Fu et al. 2012). To create the tandem dimer of yEGFP a PCR fragment was amplified with a forward primer carrying a HindIII site with a 9bp spacer (GCTGGTTTA) along with a reverse primer carrying SpeI, PstI and BamHI sites. The 1xEGFP vector and the PCR product were digested with HindIII and BamHI, then ligated. To further create the tandem trimeric yEGFP, a PCR fragment was amplified with a forward primer carrying an SpeI site with a spacer sequence as above in combination with a reverse primer carrying a PstI site. The PCR fragment and the tandem 2xyEGFP were digested with SpeI and PstI and ligated. The resulting plasmids were verified by DNA sequencing, linearized and integrated into the genome (URA3).

For FCS, whole cell extracts were prepared from 50 OD of exponentially growing cells by bead beating in buffer E. The clarified extracts were measured using Labtek chambers coated with BSA (1mg/ml) for 30 minutes and washed with PBS twice. In FCS, the changes in fluorescence intensity reflect the fluctuations of the number of particles as a function of time. Samples were diluted so that on average five molecules were present in the detection volume at a time. The average number of molecules was calculated for a serial dilution series by fitting the autocorrelation curves with a single component fitting.

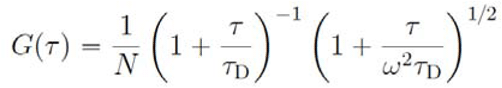

where N is the average particle number of species in the sampling volume and τ_D_ is the residence time of species within the sampling volume, with τ_D_=ω^2^ _xy_/8D with D the diffusion coefficient of the species and ω=ω_z_/ω_*xy*_ is the aspect ratio of the sampling volume. For measuring the stoichiometry of Set1C in wildtype and *sdc1* mutants in cell extracts, intensity distributions were obtained using a Zeiss LSM 780 confocal microscope in fluorescence correlation spectroscopy and photon counting histogram mode. To quantify molecular brightness, we used Fluorescence intensity distribution analysis (FIDA), which discriminates different fluorescent species according to their molecular brightness. FIDA was performed as described (Kask et al. 1999) and the experiment was repeated independently three times.

### Auxin Inducible Degron

To make strains for the auxin-inducible degradation experiment, pURA3-pAHD1-mTIR1-CYC1-pA carrying the ADH1 promoter to express OsTIR1 was integrated at the URA3 locus to obtain RC125. Then an N-terminal AID-FLAG tag was added to the N-terminus of *set1* to obtain RC126. *jhd2* was deleted in this strain to obtain RC127. Overnight cultures were inoculated into fresh YEPD (O.D. 0.1), grown to O.D. 0.4 at the respective temperatures. Cells were then arrested by adding α-factor to 15nM for 4-6 hr. To induce degradation, 3-indoleacetic acid (IAA; Sigma I2886) was added to a final concentration of 0.5mM.

### ChIP-Seq Analysis

ChIP-Seq reads were mapped to sacCer3 *S.cerevisiae* genome assembly using BWA and parallel version of pBWA (0.5.9-r32-MPI-patch2). After mapping, uniquely mapped were filtered (-q 1) and PCR duplicates (reads with same start and mapped to same strand) were removed using Samtools (rmdup)(Li and Durbin 2010). Read lengths were computationally extended to 150bp strand specifically and stored in Bam format. The Bam files were subject to coverage calculation using the bamCoverage utility of deepTools (Ramírez et al. 2014) for a sliding window of 40bp, normalized using RPKM parameters and stored in BigWig format. Additionally, these files were then subtracted by the mock IP control. Thus, all the resultant data were directly comparable because of RPKM normalization and control subtraction. Several data matrices were created for different loci of TSS +/- 1.5KB, averaged metagene from the BigWig files via computeMatrix and visualized using plotHeatmap utilities of deepTools (also referred to as high resolution intensity plots or heatmaps). GO analysis was performed using the web-based tool FuncAssociate (Berriz et al. 2003). Composite profile plots were generated using custom code in R statistical language where the input was the above computed matrices using deepTools. BigWig files were visualised as coverage tracks using the UCSC genome browser (Karolchik et al. 2008).

For qChIP experiments, reads were first aligned to sacCer3 reference and then to sPombe reference genomes (obtained via PomBase; (McDowall et al. 2015). Note that reads were independently aligned to both genomes thus allowing for common mappings as compared to left-over read alignment to reduce the mismatch bias. Percentage of reads aligned per sample per genome was documented in a tabular format.

### RNA-Seq Analysis

RNA-seq data were processed mostly using Tuxedo suite and custom scripts for further downstream analysis (Trapnell et al. 2012). Tuxedo suite consists of TopHat (aligns the reads and map them to the genome) (Trapnell et al. 2009), Cufflinks (uses the read alignment map to assemble reads into transcripts) (Trapnell et al. 2010), Cuffdiff (takes aligned reads from multiple conditions and performs differential gene expression analysis) (Trapnell et al. 2013). The counts are calculated and recorded in an expression unit of ‘Fragments Per Kilobase of exon model per Million mapped reads’ (RPKM). We used a cutoff of FPKM>4 for a gene to be called as expressed. Plots were generated using cummeRbund and custom scripts written in R statistical language.

### Data availability

The ChIP-seq and RNA-seq data used in this publication have been deposited to the [name of the database] database [URL] and assigned the identifier [accession].

## Supporting information

## Acknowledgements

Thanks to Dani Fujimori for discussions and Vincent Geli for providing the Myc tagged version of Set1. Thanks also to the Dresden Genome Center led by Andreas Dahl for ChIP-seq and RNA-seq Ilumina sequencing and Andreas Petzold for initial data handing.

## Author contributions

RC performed all the experiments except the massively parallel DNA sequencing and Fig 1F which was performed with SA. The computational analyses were performed by SS with additional contributions by RC and AFS. The project was based on initial results from AR. The experiments were planned and the manuscript written by RC and AFS.

## Competing Interest

The authors declare no competing interests.

