## Supplementary material for "The Set1 complex is dimeric and acts with Jhd2 demethylation to convey symmetrical H3K4 trimethylation"

**Supplementary Material**
**9 Supplementary Figures**
**1 Supplementary Table – an Excel file not included in this document**
**Extended Materials and Methods**

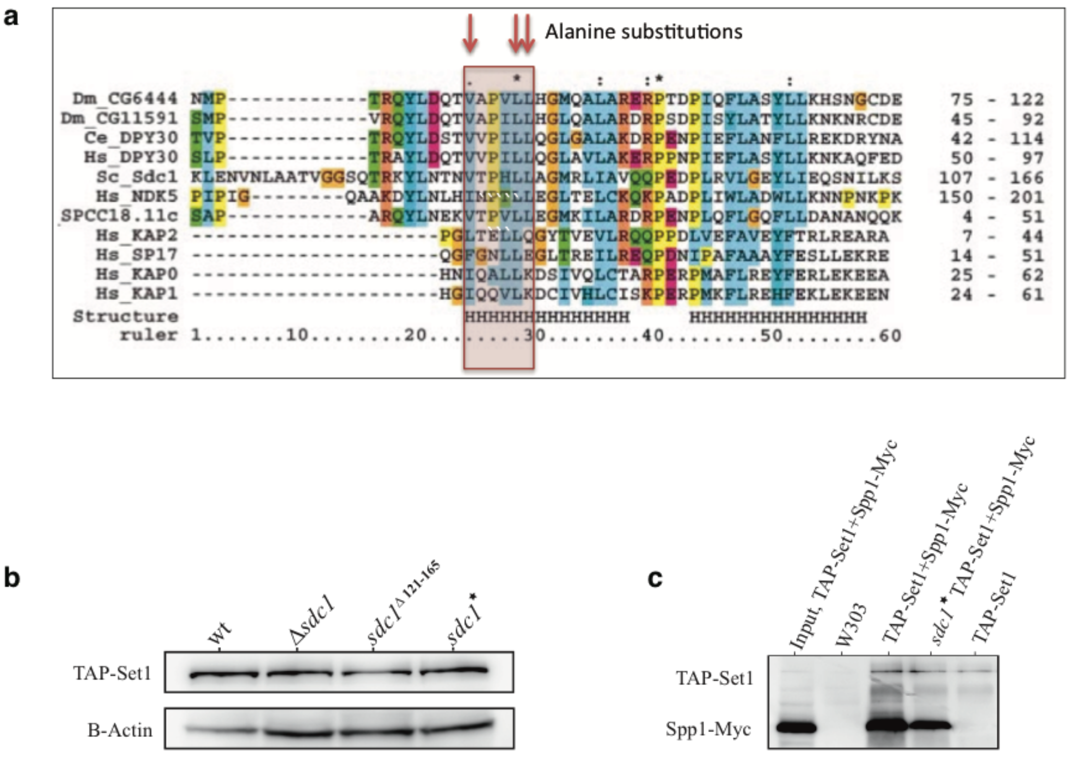

**Supplementary Figure 1. Sdc1 mediated dimerization of yeast Set1 complex.**

**(A)** Multiple sequence alignment of the dimerizaton region of the protein kinase A regulatory subunit II with Dpy30 homologues and other proteins (from Roguev et al, 2001). Arrows indicate the three amino acids that were changed to alanine in *sdc1*.*

**(B)** Expression levels of TAP-Set1 protein in wild type and *sdc1* mutant strains evaluated by Westerns using whole cell extracts and beta-actin (B-actin) as loading control.

**(C)** Spp1 associates with the monomeric Set1 complex. Spp1 was tagged with a Myc epitope in wild type and sdc1* strains that carried TAP-Set1. TAP-Set1 was immunoprecipitated from whole cell extracts and evaluated by Western with an anti-myc antibody. Input, whole cell extract from the *TAP-set1; spp1-myc* strain. W303, whole cell extract from the W303 parental strain.

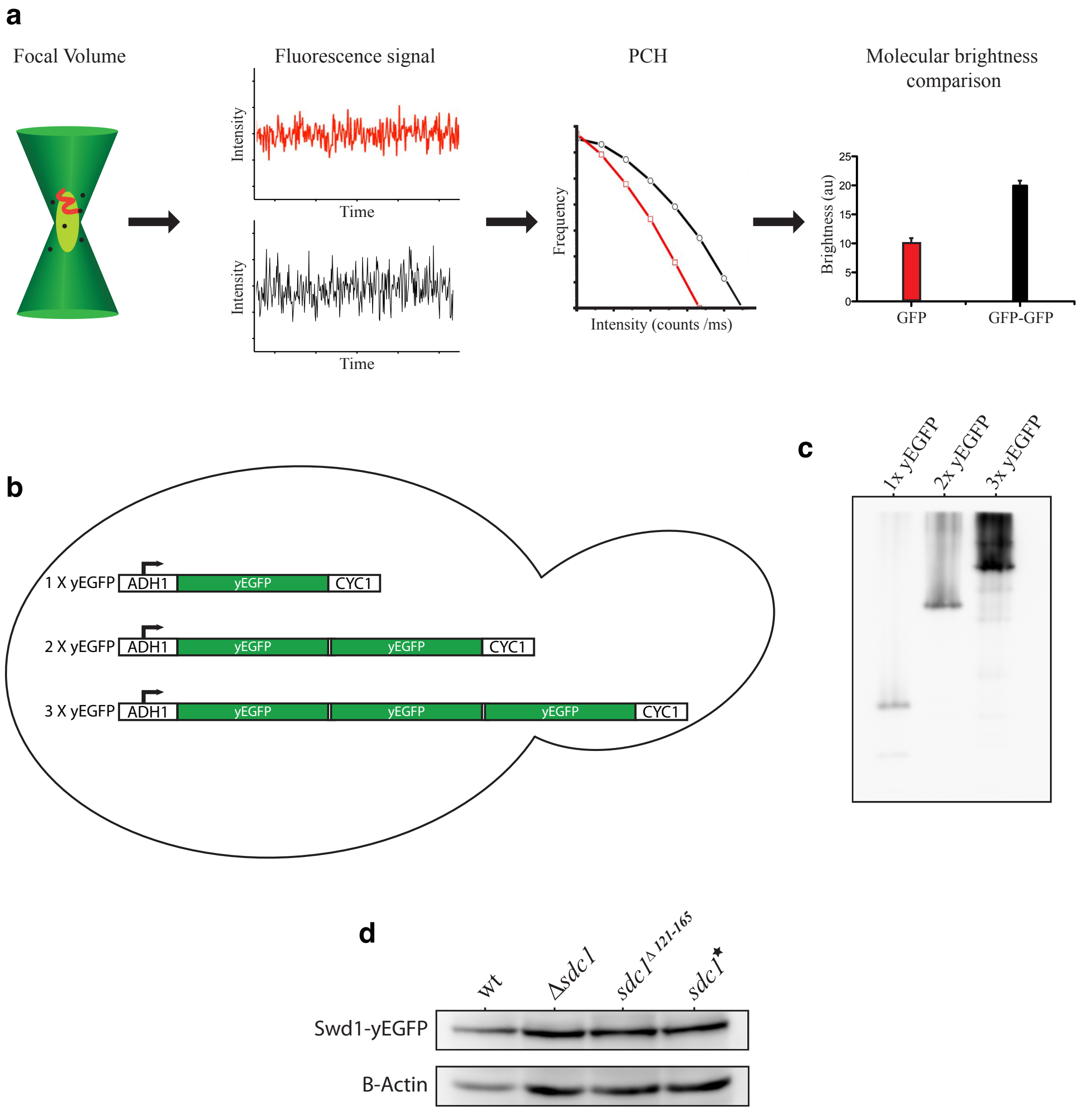

**Supplementary Figure 2. Molecular analysis of Sdc1 mediated dimerization of Set1C by Fluorescence Correlation Spectroscopy (FCS).**

**(A)** Outline of the FCS experiment. Fluorescence fluctuation data from whole cell extracts were analyzed by Photon Counting Histogram (PCH) analysis to estimate the number of molecules and stoichiometry. A PCH analysis of GFP (open red square) and tandem GFP-GFP (open black circle) construct followed by molecular brightness analysis reveals the difference between single and tandem dimer GFPs.

**(B)** Schematic representation of yeast expressing yEGFP (yeast codon optimized EGFP) transgenes as illustrated integrated at the *ura3* locus.

**(C)** Whole cell extracts of yeast expressing 1x, 2x and 3x yEGFP were analyzed by immunoblotting for GFP.

**(D)** Expression levels of Swd1-yEGFP protein in wild type and *sdc1* mutant strains. Whole cell extracts were resolved by SDS-PAGE and analysed by immunoblotting for the GFP tag with beta-actin as loading control.

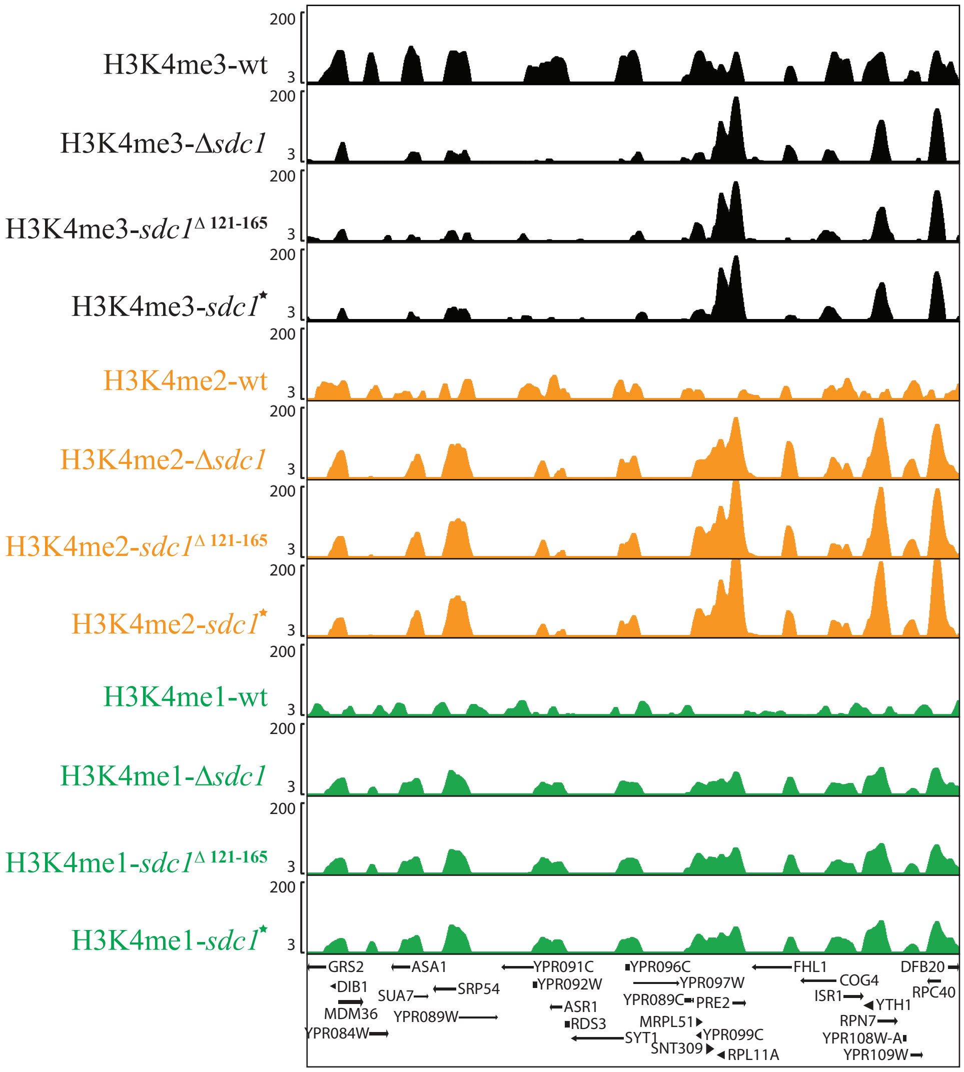

**Supplementary Figure 3. Altered distribution of H3k4 methylation by monomeric Set1C.**

Screen shot of H3K4me3 (top, black). H3K4me2 (middle, orange) and H3K4me1 (green, bottom) ChIP-seq from wt, *Δsdc1, sdc1*$^{\Delta 121\text{-}165}$ and *sdc1** strains as indicated with gene diagrams below.

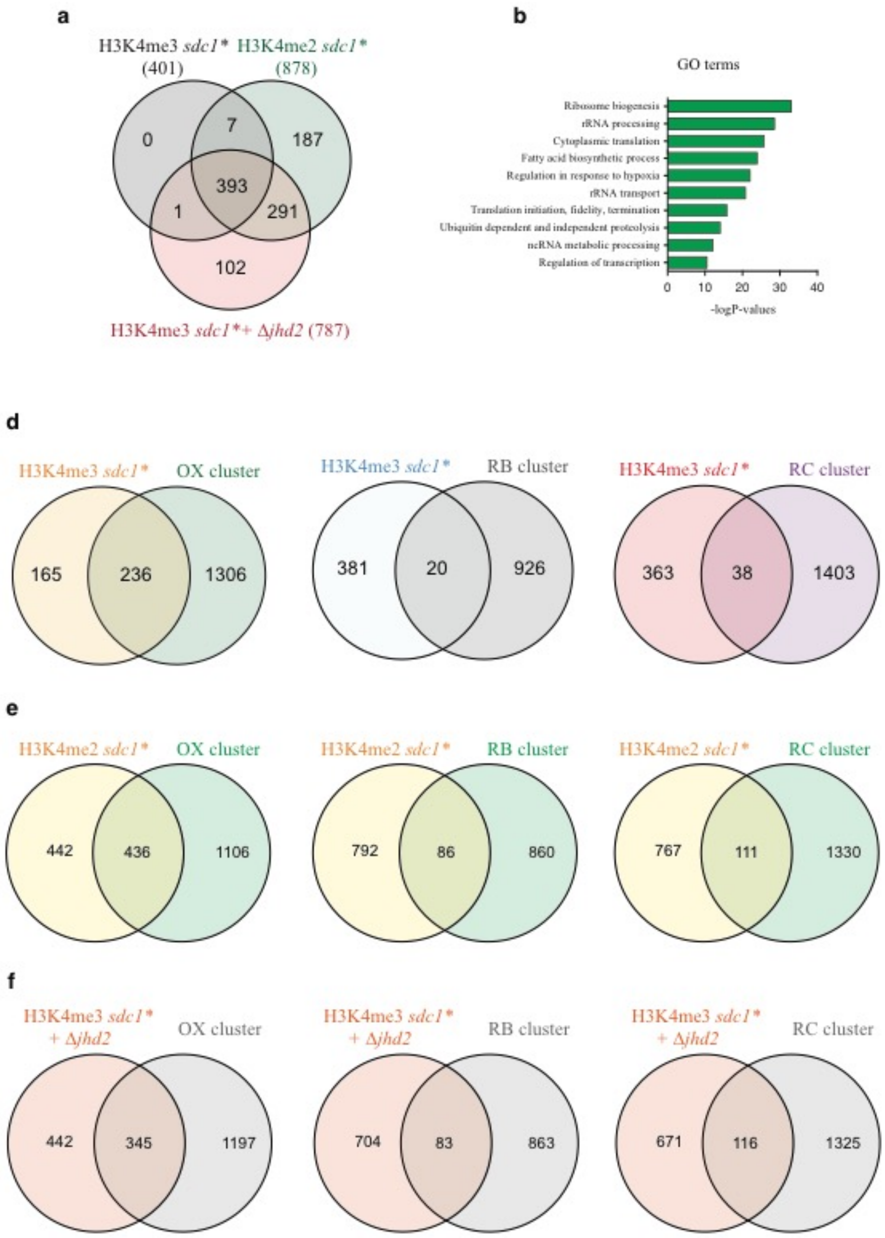

**Supplementary Figure 4. Relationships between the cluster 1s.**

**(A)** Venn diagram of the relationship between the H3K4me3 *sdc1** cluster 1 (401), the *sdc1** H3K4me2 cluster 1 (878) and the *sdc1*/Δjhd2* H3K4me3 cluster 1 (787).

**(B)** GO term summary for the H3K4me3 *sdc1** cluster 1 (401). GO analysis employed yeastract (REF).

**(C)** Venn diagrams of the relationships between H3K4me3 sdc1* (401; cluster 1) and the 1542 oxidative (OX), 946 reductive building (RB) and 1441 reductive charging (RC) genes in the YMC[1].

**(D)** same as (c) except for the H4K4me2 *sdc1** (878; cluster 1).

**(E)** same as (c) except for the H3K4me3 *sdc1*/Δjhd2* (787; cluster 1).

a

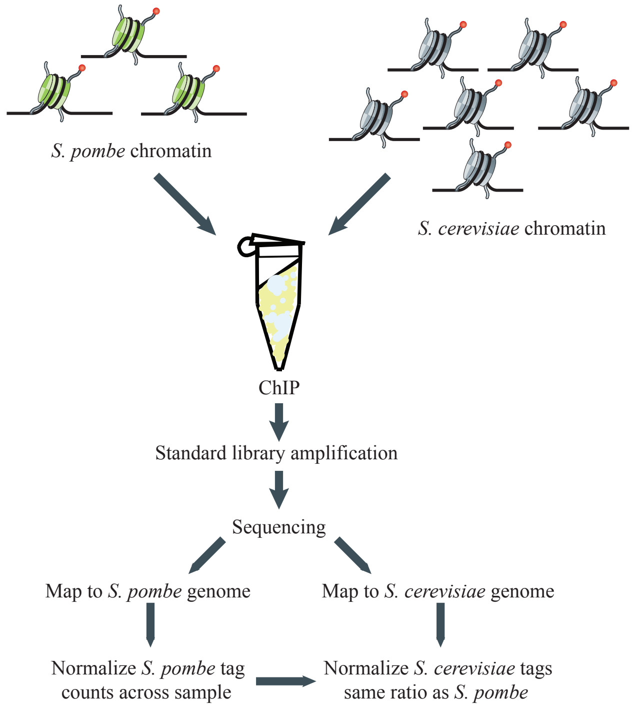

b

| Sample | *S. cerevisiae* | *S. pombe* | Adj. Factor | | Ave. Adj. Factor |
|---|---|---|---|---|---|
| H3K4me3_clone 1_*wt*_DS1 | 48.33 | 8.59 | 2.81 | | |
| H3K4me3_clone 2_*wt*_DS1 | 71.31 | 16.29 | 2.18 | H3K4me3 | |
| H3K4me3_clone 1_*wt*_AR42 | 74.79 | 15.39 | 2.42 | *wt* | 2.43 |
| H3K4me3_clone 2_*wt*_AR42 | 75.01 | 16.22 | 2.31 | | |
| H3K4me3_clone 1_*sdc1** | 31.97 | 44.6 | 6.79 | *Sdc1** | 8.58 |
| H3K4me3_clone 2_*sdc1** | 26.82 | 57.06 | 10.36 | | |
| H3K4me3_clone 1_*Δjhd2* | 76.57 | 13.74 | 0.87 | *Δjhd2* | 0.89 |
| H3K4me3_clone 2_*Δjhd2* | 70.46 | 13.27 | 0.91 | | |
| H3K4me3_clone 1_*sdc1**_*Δjhd2*_AR42 | 26.41 | 26.41 | 3.71 | *sdc1**_*Δjhd2* | 4.34 |
| H3K4me3_clone 2_*sdc1**_*Δjhd2*_AR42 | 42.23 | 43.08 | 4.96 | | |
| H3K4me2_clone 1_*wt*_DS1 | 71.92 | 15.71 | 2.28 | | |
| H3K4me2_clone 2_*wt*_DS1 | 68.76 | 14.75 | 2.33 | H3K4me2 | |
| H3K4me2_clone 1_*wt*_AR42 | 57.21 | 57.21 | 3.71 | *wt* | 2.97 |
| H3K4me2_clone 2_*wt*_AR42 | 69.16 | 69.16 | 3.56 | | |
| H3K4me2_clone 1_*sdc1** | 42.18 | 21.35 | 3.01 | *Sdc1** | 3.3 |
| H3K4me2_clone 2_*sdc1** | 38.01 | 22.96 | 3.59 | | |
| H3K4me2_clone 1_*Δjhd2* | 65.52 | 65.52 | 1.02 | *Δjhd2* | 1.01 |
| H3K4me2_clone 2_*Δjhd2* | 59.27 | 59.27 | 1.01 | | |
| H3K4me2_clone 1_*sdc1**_*Δjhd2*_AR42 | 68.74 | 68.74 | 0.91 | *sdc1**_*Δjhd2* | 1.1 |
| H3K4me2_clone 2_*sdc1**_*Δjhd2*_AR42 | 76.03 | 14.74 | 1.3 | | |
| H3K4me1_clone 1_*wt*_DS1 | 56.84 | 23.59 | 1.2 | | |
| H3K4me1_clone 2_*wt*_DS1 | 56.18 | 23.45 | 1,19 | H3K4me1 | |
| H3K4me1_clone 1_*wt*_AR42 | 42.11 | 42.11 | 1.75 | *wt* | 1.44 |
| H3K4me1_clone 2_*wt*_AR42 | 63.34 | 63.34 | 1.63 | | |
| H3K4me1_clone 1_*sdc1** | 70.25 | 70.25 | 0.5 | *Sdc1** | 0.59 |
| H3K4me1_clone 2_*sdc1** | 67.91 | 16.29 | 0.69 | | |
| H3K4me1_clone 1_*Δjhd2* | 57.46 | 16.36 | 0.82 | *Δjhd2* | 0.85 |
| H3K4me1_clone 2_*Δjhd2* | 58.97 | 18.16 | 0.89 | | |
| H3K4me1_clone 1_*sdc1**_*Δjhd2*_AR42 | 78.24 | 78.24 | 0.25 | *sdc1**_*Δjhd2* | 0.3 |
| H3K4me1_clone 2_*sdc1**_*Δjhd2*_AR42 | 75.92 | 75.92 | 0.36 | | |

**Supplementary Figure 5. Quantifying changes in histone modification using spike-in ChIP-sequencing.**

**(A)** Schematic representation of ChIP-sequencing protocol using 1/3$^{rd}$ input control of wild type *S. pombe* chromatin. The *S. cerevisiae* and *S. pombe* cells were mixed after formaldehyde fixation and then further processed as one sample. After sequencing and mapping, total sequence tags to the two genomes were used to generate adjustment factors. **(B)** Table presenting the spike-in results and adjustment factors. The sample column lists the strain used. The experiments were performed in duplicate using a 2:1 *S. cerevisiae:S. pombe* input ratio mixed together after neutralizing the crosslink. The columns '*S. cerevisiae'* and '*S. pombe'* show the percentage of unique reads mapped to that genome for each sample. These figures do not total 100% due to non-unique and non-yeast reads. The adjustment factor for each sample is the difference between the expected 2:1 ratio and the result. This was then averaged for each duplicate. Except for the wt average adjustment factor, which is simply the average from the wt samples, the average adjustment factor in the far right column was calculated by multiplying the average *S. pombe* percentage yield with the wt average adjustment factor and then divided by the *S. cerevisiae* percentage yield. For example, *S. pombe* H3K4me3 ChIP-seq reads were strongly over represented when mixed with *sdc1** chromatin but not when mixed with *Δjhd2* chromatin. Thereby indicating that H3K4me3 was greatly reduced for *sdc1** but not *Δjhd2* chromatin and giving average adjustment factors of 8.58 and 0.89 respectively.

**a**

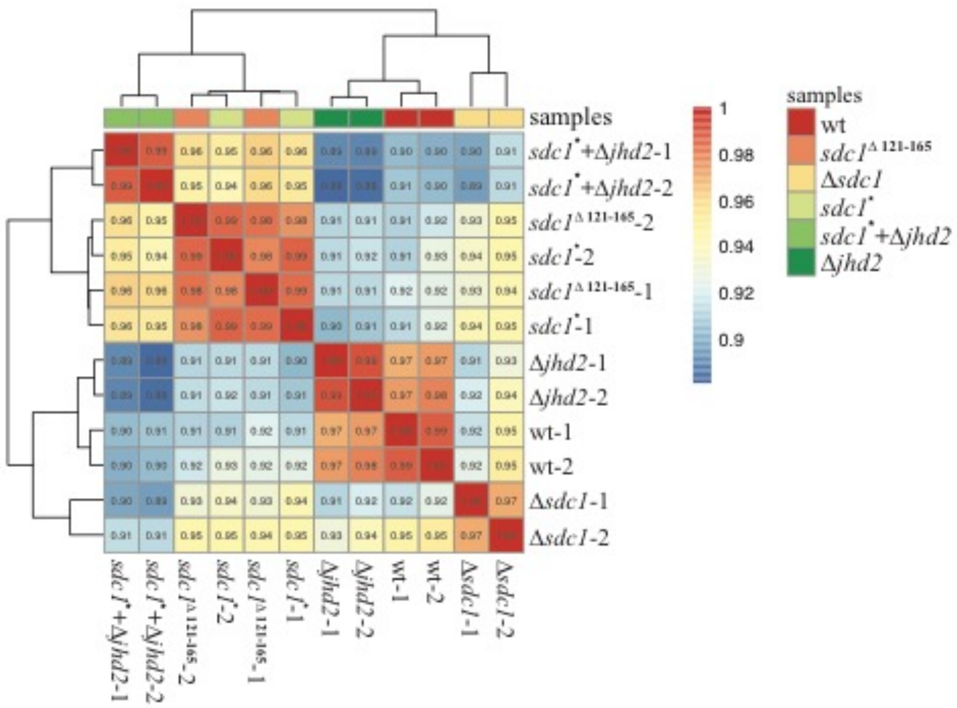
samples
sdc1*+Δjhd2-1
sdc1*+Δjhd2-2
sdc1Δ 121-165-2
sdc1*-2
sdc1Δ 121-165-1
sdc1*-1
Δjhd2-1
Δjhd2-2
wt-1
wt-2
Δsdc1-1
Δsdc1-2
1
0.98
0.96
0.94
0.92
0.9
samples
wt
sdc1Δ 121-165
Δsdc1
sdc1*
sdc1*+Δjhd2
Δjhd2

**b**

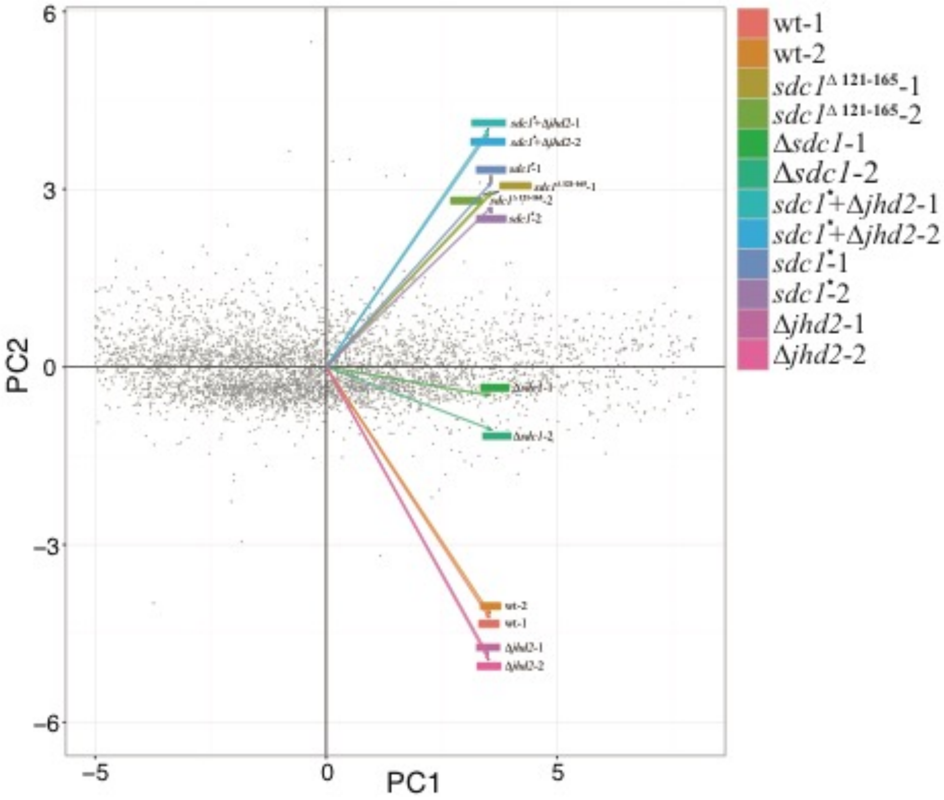
PC2
PC1
6
3
0
-3
-6
-5
0
5
wt-1
wt-2
sdc1Δ 121-165-1
sdc1Δ 121-165-2
Δsdc1-1
Δsdc1-2
sdc1*+Δjhd2-1
sdc1*+Δjhd2-2
sdc1*-1
sdc1*-2
Δjhd2-1
Δjhd2-2

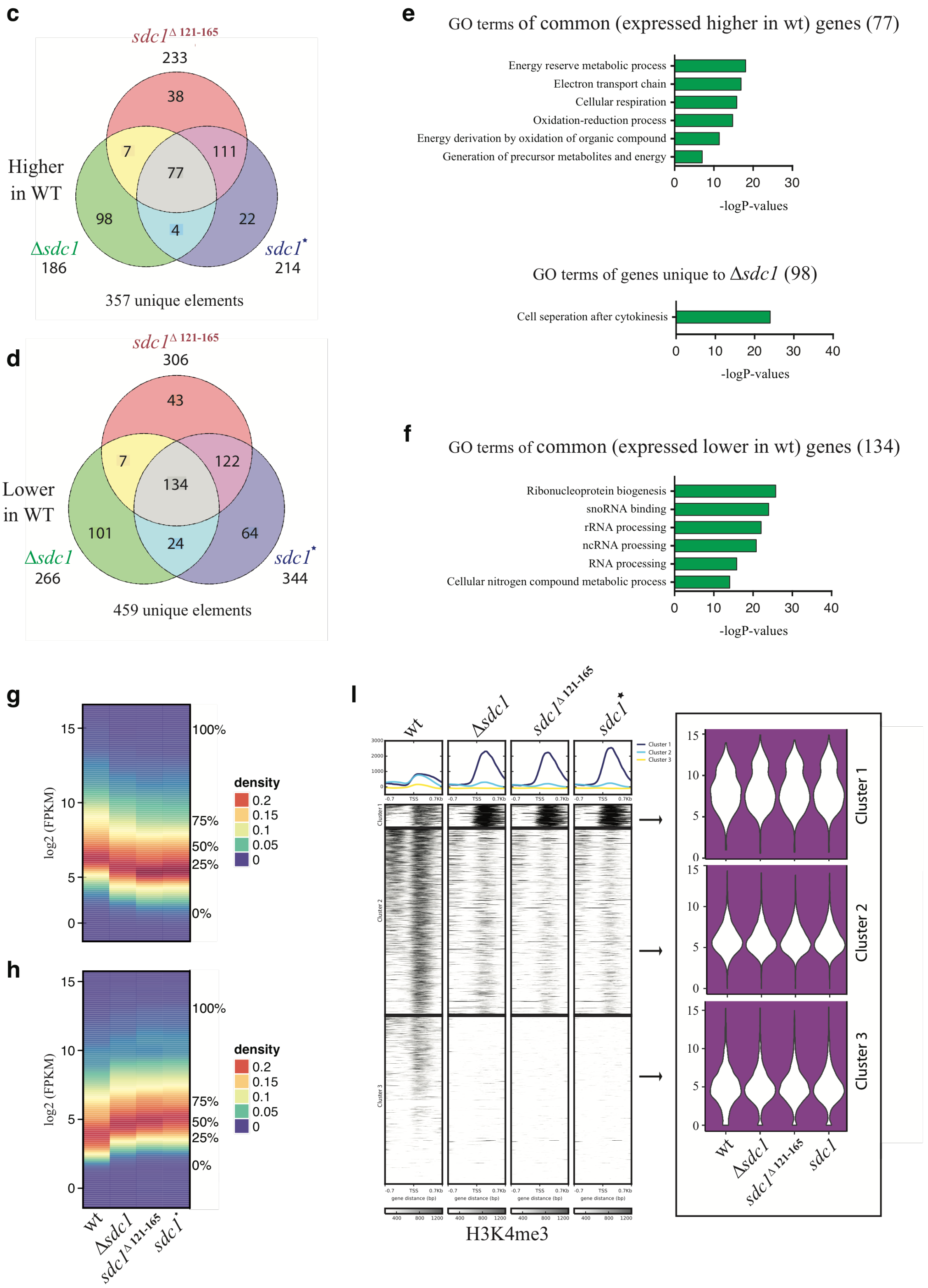

**Supplementary Figure 6. Monomeric Set1 complex has a moderate effect on gene expression compared to wild type.**

Duplicates of total poly A plus RNA for wt and the mutant strains as indicated were Illumina 2500 paired-end sequenced. After removal of PCR duplicates and non-mappable reads, the data sets were equated to ~10 million reads each for further analyses.

**(A)** Heatmap plot showing gene expression correlations between wt and the mutant strains based on their quantitative differential gene expression analysis revealed that wt and *sdc1* mutants were closely related. *sdc1*$^{\Delta 121\text{-}165}$ and *sdc1** were highly similar whereas *Δsdc1* was slightly more related to wt whereas *Δjhd2* had almost no effect when either removed from wt or *sdc1** strains.

**(B)** Principal Component Analysis (PCA) plot showing the same effect as observed with the heatmap in (a). Again *sdc1*$^{\Delta 121\text{-}165}$ and *sdc1** cluster together whereas *Δsdc1* was closer to wt.

**(C)** Venn diagram showing the overlap of mRNAs downregulated more than two fold compared to wt between the three *sdc1* mutants. In total, 357 genes were encompassed, of which only 77 (22%) were common to all three duplicate experiments. Again note the concordance between *sdc1*$^{\Delta 121\text{-}165}$ and *sdc1** (188/233; 81% and 188/214; 88%) compared to *Δsdc1* (84/186;45% and 81/186; 44%).

**(D)** As for (c) except for mRNAs upregulated more than two-fold compared to wt. Of the total 459 genes, 134 (29%) were common to all three duplicate experiments. Concordance - *sdc1*$^{\Delta 121\text{-}165}$ and *sdc1** (256/306; 84% and 256/344; 74%); *Δsdc1* (158/266; 59% and 141/266; 53%).

**(E)** Gene Ontology enrichment analysis of the 77 overlapping genes in (c) showing the top 6 terms, as well as the 98 genes unique to *Δsdc1.* In this group of 98, the top GO term was cell separation after cytokinesis. This is notable because we noticed that the *Δsdc1* strain tends to clump displaying signs of defective cell division. This is not apparent with the other two mutant *sdc1* strains, suggesting that Sdc1 conveys a function related to the expression of genes involved in cytokinesis in addition to the functions of the DPY-30 box, which includes the dimerization and Bre2 interactions.

**(F)** Gene Ontology enrichment analysis the of the 134 overlapping genes in (d) showing the top 6 terms.

**(G)** A quantification matrix plot displaying the 357 mRNAs of (c) that were downregulated by two-fold or more in the *sdc1* mutant strains.

**(H)** Same as (e) except displaying the 459 mRNAs of (d) that were downregulated more than two-fold in the *sdc1* mutant strains.

**(I)** Violin plots of gene expression values from the promoters in each of the three clusters of Figure 3b, part of which is reproduced here for visual clarity. Whereas the dramatic changes in H3K4me3 are obvious, no corresponding alterations in gene expression can be observed.

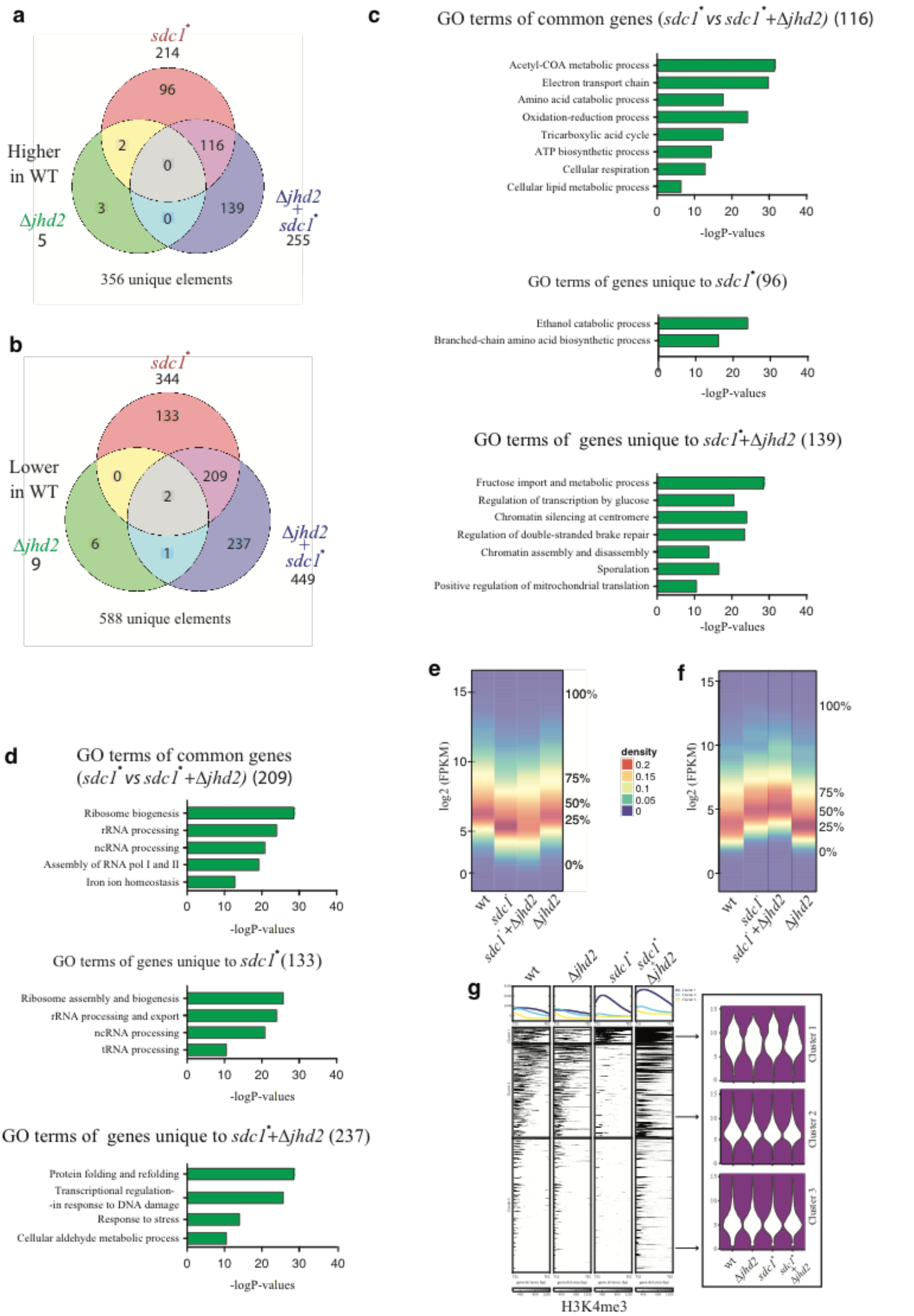

**Supplementary Figure 7. Deletion of *jhd2* has virtually no effect on mRNA expression. (A)** Venn diagram showing the overlap of mRNAs downregulated more than two fold compared to wt between the three mutant strains. In total, 356 genes were encompassed with only 5 affected by the loss of Jhd2. Excluding the 3 unique to *Δjhd2*, only 32% (116/353) were common to *sdc1** and *sdc1**+*Δjhd2*.

**(B)** As for (a) except for mRNAs upregulated more than two-fold compared to wt. Of the total 582 genes (excluding 6 unique to *Δjhd2)*, 36% (209/582) were common to *sdc1** and *sdc1**+*Δjhd2*.

**(C)** Gene Ontology enrichment analyses of the genes in (a); at top the 116 overlapping genes; in the middle the 96 genes unique to *sdc1**; below the 139 genes unique to *sdc1**+ *Δjhd2.*

**(D)** Gene Ontology enrichment analyses of the genes in (b); at top the 209 overlapping genes; in the middle the 133 genes unique to *sdc1**; below the 237 genes unique to *sdc1**+ *Δjhd2.*

**(E)** A quantification matrix plot displaying the 356 mRNAs from (a) downregulated more than two-fold in the *sdc1* mutant strains.

**(F)** Same as (e) except the 588 mRNAs from (b) upregulated more than two-fold.

**(G)** Violin plots of gene expression values from the promoters in each of the three clusters of Figure 6a, part of which is reproduced here for visual clarity. Whereas the dramatic changes in H3K4me3 are obvious, no corresponding alterations in gene expression can be observed.

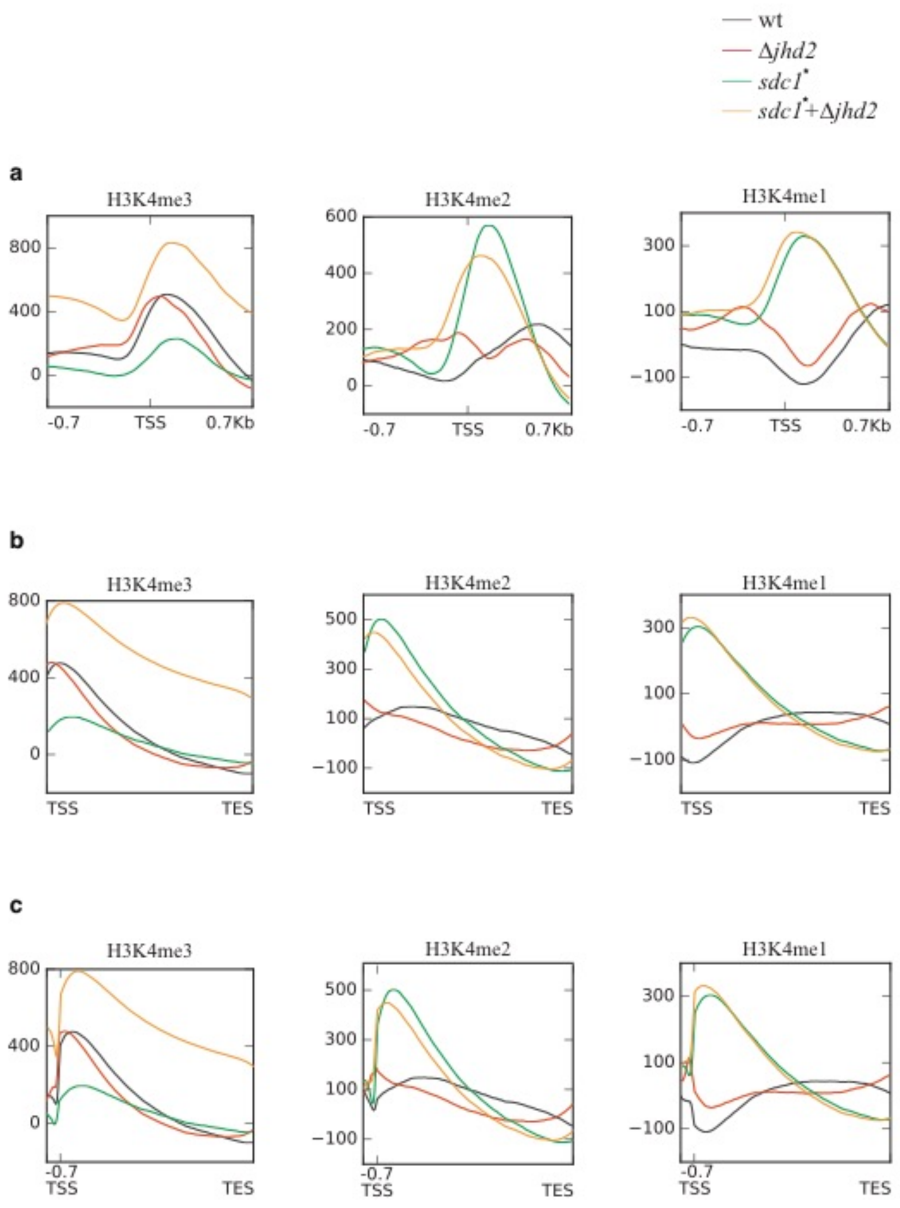

**Supplementary Figure 8. Role of Jhd2 on appropriate distribution of H3K4 methylation across genome.**

**(A)** Metagene plots showing the average distribution of H3K4me3, me2 and me1 centered on the transcription start site (TSS +/- 0.7kb). The normalized distribution of the methylation marks on an average gene is shown for WT (black line), Δ*jhd2* deletion (red line), *sdc1** (green line) and the combination *sdc1** + Δ*jhd2* (orange line).

**(B)** The same as (a) except from transcription start site (TSS) to transcription end site (TES).

**(C)** The same as (a) and (b) except from -0.7kb upstream of TSS to TES.

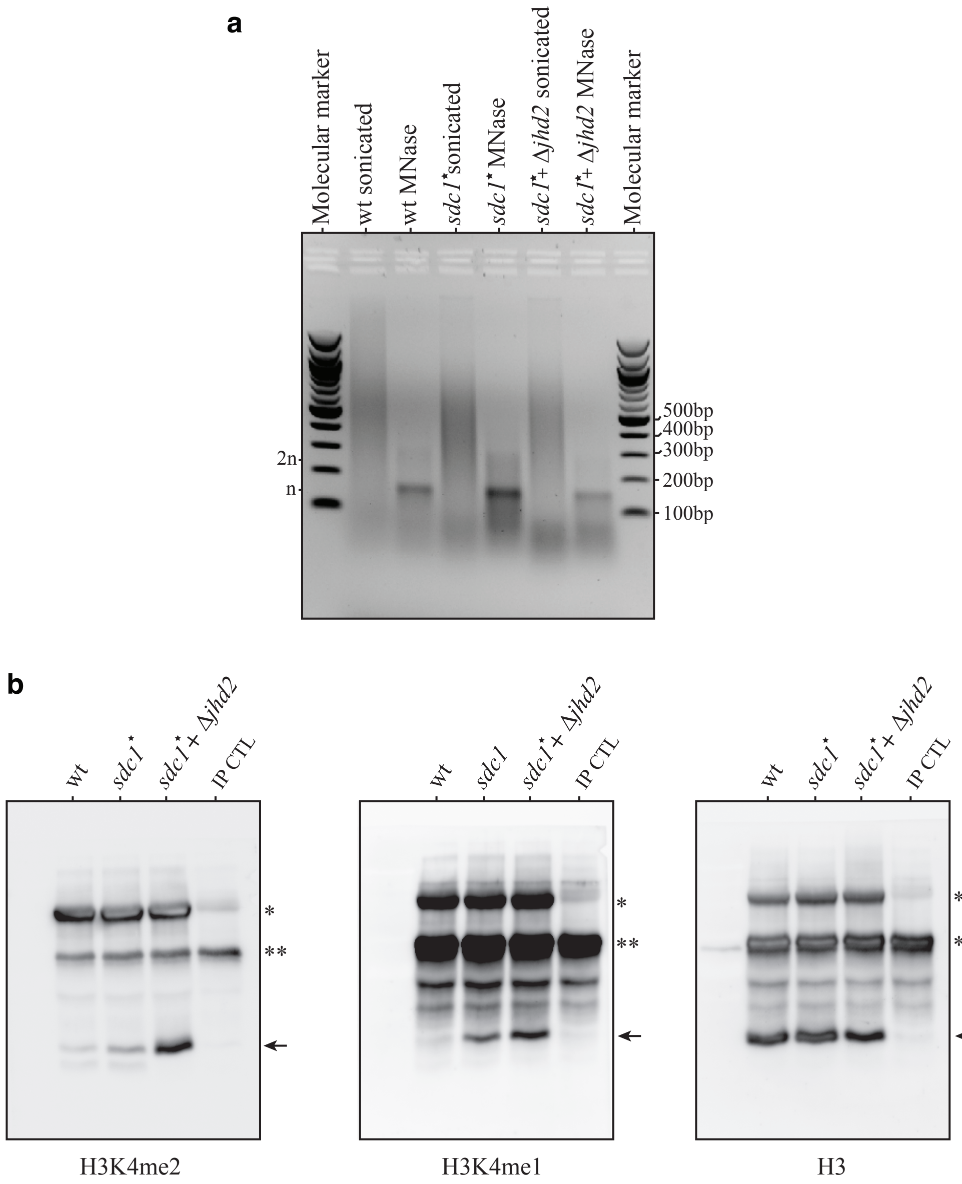

**Supplementary Figure 9. MNase digestion of yeast chromatin to mononucleosomes**

Sonicated formaldehyde crosslinked, ChIP-ready, chromatin was digested with micrococcal nuclease (MNase). Aliquots were withdrawn, de-crosslinked and DNA was fractionated on a 2% 1xTBE agarose gel and visualized with EtBr. M, markers (NEB 100bp DNA ladder); n, 2n mononucleosomes, dinucleosomes.

(**A**) Samples from the indicated strains before (sonicated) or after MNase digestion.

(**B**) Full gel images of Figure 6E

### Extended Materials and Methods

#### Yeast Strains

Strains used in this study are listed in Supplementary Table I. All haploid strains were derived from MGD353-13D [2]. Yeast transformations were performed as described [3]. Gene disruptions and gene tagging were performed as described [2,4]. Correct cassette integrations were confirmed by PCR and western blot analysis (for tagging) or PCR and genomic Southern blot (for disruptions).

**Protein assays and immunoblotting**

For size exclusion chromatography, a 10/30 Superose 6 size exclusion column (HR, Pharmacia) was loaded with 500μl of cleared crude cell extract from a TAP-tagged strain and run in glycerol-free buffer E (20 mM HEPES-NaOH, pH 8.0, 350 mM NaCl, 0.1% Tween 20, 10% glycerol, 1ug each of leupeptin, aprotinin, and pepstatin A, 1 mM phenylmethylsulfonyl fluoride;[5]). Fractions were resolved on an 8% SDS-PAGE gel and analyzed by immunoblotting with peroxidase-anti-peroxidase (PAP, Sigma) diluted 1:1000, for detection of the protein A region within the TAP tag using the ECL kit (Amersham Pharmacia Biotech). For co-immunoprecipitation of TAP-Set1 and Myc-Set1, 50ml yeast cultures were grown under selection to mid-log phase. Whole cell extracts were prepared by bead beating in 500μl of lysis buffer E and protein levels were normalized using a Bradford assay. A portion of each clarified lysate was reserved for input. For TAP-Set1 immunoprecipitations, the remaining lysates were incubated with 20μl of IgG-Sepharose (Pharmacia), equilibrated in buffer E added to normalized extracts and incubated for 2h at 4°C with rotation. The immunoprecipitated samples were washed three times in 1ml of lysis buffer. The remaining beads from the final wash were resuspended in 2xSDS sample buffer with 10% β-mercaptoethanol and boiled. Each immunoprecipitated sample was split in half and 10μl was loaded per lane on a 10% SDS polyacrylamide gel, transferred to PVDF membrane and probed using α-Myc antibody (Myc; 9E10, catalog number 11667203001, Roche Applied Science) or PAP antibody (catalog number P1291, Sigma). For immunoprecipitation of Myc-Set1, the above procedure was performed with the following modifications. To each sample, 2μg of α-Myc antibody (9E10, catalog number 11667203001, Roche Applied Science) was added to normalized extracts and incubated at 4°C for 2h. After 2h, 20μl of Protein G-Sepharose (catalog number 17-0618-01, GE Healthcare) was added to each sample and incubated for 1h at 4°C. The immunoprecipitated samples were washed three times with 1ml of buffer E and the remaining beads were resuspended in 20μl of 2xSDS sample buffer with 10% β-mercaptoethanol and boiled. Samples were analysed on an 10% SDS polyacrylamide gel, transferred to a PVDF membrane and probed with PAP antibody (PAP; catalog number P1291, Sigma). For co-immunoprecipitation studies of TAP-Set1 with Pol II CTD, the cell extracts were prepared as above except that the clarified whole cell extract was supplemented with 2mM MgCl2 and 2μl Benzonase (Novagen, 71205) followed by incubation at room temperature for 10 min. Histones were purified using a protocol adapted from [6,7]. Briefly, harvested yeast pellets from 2 liters log-phase cultures were homogenized using a bead beater (BioSpec) in a modified Nuclear Isolation Buffer (0.25M Sucrose, 60mM KCl, 15mM NaCl, 5mM MgCl2, 1mM CaCl2, 20mM

HEPES pH8.0, 0.5mM spermine, 2.5mM spermidine, 0.8% Triton X-100, 10mM Na butyrate and protease inhibitors). The homogenate was centrifuged at 32,000g for 15 minutes, the crude chromatin pellet resuspended in 0.25N HCl, sonicated and rotated at 4°C for one hour. Acid insoluble material was cleared by centrifugation and discarded. Acid soluble material was purified in batch using BioRex70 ion exchange resin (BioRad). Samples were dialyzed against HPLC buffer A ($H_2O$ with 0.1% TFA) and separated by C4 RP-HPLC over multi-step acetonitrile gradients. Histone containing fractions from the C4 separation were collected, lyophilized, and boiled in 1xSDS sample buffer with 10% β-mercaptoethanol. Samples were analysed on an 12% SDS polyacrylamide gel, transferred to a PVDF membrane and immunoblotted with antibodies against H3K4me1 (pAB-037-050, 1:2000, Diagenode), H3K4me2 (pAB-035-050, 1:1000, Diagenode), H3K4me3 (ab8580, 1:1000, Abcam), and H3 (ab1791, 1:2000, Abcam) in TBST+5% skim milk at 4°C. Blots were then washed twice with PBST (PBS, 1% Tween20), incubated with secondary antibody conjugated to HRP (cat no. 3140, 1:10000, Thermo), and exposed to Enhanced Chemiluminescence reagents (Pierce) and imaged using ImageQuant LAS400 imaging System (GE). The blots were quantified using ImageQuantTL image analysis program (GE).

**ChIP-seq**

All ChIP-seq experiments were performed in duplicates. Exponentially growing cells ($2\times10^8$) were collected and washed with water at room temperature (RT) and resuspended in 40ml of PBS (RT) containing 1% paraformaldehyde (in a fume-hood) for fixation. The cultures were agitated slowly for 20 min at room temperature. The cell fixation quenched by adding 2.5ml of 2.5M glycine (drop wise) and continued to agitate for 5 min at room temperature before centrifugation at 3000rpm in the cold room with further steps on ice. The cells were washed in 2x25ml ice cold PBS. The cell pellet was suspended in 400μl ice cold lysis buffer (50mM HEPES-KOH pH 7.5, 150mM NaCl, 1mM EDTA, 1% Triton X-100, 0.1% sodium deoxycholate, 0.1% SDS) supplemented with complete protease inhibitor cocktail (Roche, Ref. 04693116001). The cell suspension was then transferred to a precooled Sarstedt tube with one volume acid-washed glass beads (0.5mm glass beads, Biospec). Cells were lysed using BioSpec Mini-Bead Beater until > 95% were lysed, as evaluated by microscopy before centrifugation at 4000g. The chromatin was gently aspirated using a blue tip and transfered to a glass sonication tube (TC16, LGC Genomics, KBS-0059-545). The volume of the chromatin mix was made up to 800μl by adding lysis buffer. The chromatin was then sonicated using Covaris S2 focused ultrasonicator (Intensity 4, 20% duty cycle, 200 cycles per burst, 10 min, at 4°C). The sonicated DNA ranged from 200-500 bp as evaluated by treating 50μl aliquots with proteinase K treatment at 65°C for 2 hr, DNA purification on QIAquick Spin Columns (QIAGEN) and electrophoresis on 2% agarose gels. Otherwise chromatin was centrifuged for 10 min at 14,000g at 4°C. The supernatant was transferred to a fresh 1.5 ml tube on ice and the protein concentration determined by Bradford (Pierce, 23246). 200μl (50μg) of the chromatin per sample was mixed with 2μg of anti-H3K4Me1, -H3K4me2, -H3K4me3, -H3 or nothing. The samples were incubated with gentle rotation on a spinning wheel for 2hr at 4°C. Protein A/G PLUS-agarose beads (Santa Cruz Biotech, sc-2003) equilibrated in lysis buffer and pre-blocked (in lysis buffer with 1.5% fish gelatin, 200μg/ml salmon sonicated sperm DNA, Staratagene, Cat no. 201190-81) for 1hr in cold room. 100μl of agarose beads was added per tube and incubated for additional 1 hour at 4°C. The beads were washed under stringent buffer conditions. First washed with 500μl cold lysis buffer once in the cold room followed by 2 times 500μl lysis buffer (RT), 15 min each on a slow spinning wheel. The beads were then twice washed for 15 minutes with 500μl of deoxycholate buffer (10mM Tris-HCl, 1mM EDTA, 0.5% sodium deoxycholate, 0.5% NP-40, 0.25M LiCl) followed by one more wash with 500μl TE (10mM Tris-HCl pH8, 1mM EDTA) for 15min. The immunoprecipitated chromatin (ChIP) was eluted in 100μl with TES buffer (50mM Tris-HCl pH8, 1.5% SDS, 10mM EDTA) at 65°C with gentle shaking for at least 15 min, twice. The two elutions were pooled and incubated at 65°C overnight to reverse the cross-linking. The samples were treated with 20μg of proteinase K in 50μl of TE pH8 and incubated at 56°C for 2 hours. The immunoprecipitated DNA and the input DNA were purified with QIAquick Spin Columns and used for library construction with Illumina's TruSeq DNA Library preparation kit. DNA libraries were validated with a 2100 Bioanalyzer (Agilent Technologies) and sequenced with NextSeq 500 System (Illumina) using default Illumina standards for base calling and read filtering [8].

For the spike-in experiments [9], after crosslinking 15 OD of *S. pombe* cells were mixed with 30 OD of *S. cerevisiae* cells immediately before cooling and bead beating.

**RNA-seq**

Three independent cultures for each strain were grown to mid-log phase and $5\times10^7$ cells collected by centrifugation at 3000g for 5 min and washed with nuclease free water. Total RNA was isolated using YeaStar RNA Kit (Zymo research, R1002). To eliminate residual DNA total RNA was treated with DNase I (RNase_free DNase, Qiagen) before purification using RNeasy Kit (Qiagen). The quality of RNA was determined using Bioanalyzer (Agilent) to verify an RNA integrity number (RIN) for each sample in the range 7-8.1. The NEBNext rRNA Depletion kit was used to remove ribosomal RNA from total RNA sample. Library construction was performed using the Illumina TruSeq RNA Sample Preparation Kit version 2 and sequenced using the Illumina Hiseq2000 sequencer.

**MNase-IP**

Samples were treated in the same way as for ChIP except after sonication 5mM $MgCl_2$, 1mM $CaCl_2$ and 2μl Micrococcal Nuclease (NEB, M0247S) were added and incubated at 37°C for 20 min [10]. The digestion was halted by shifting the reaction to 4°C and adding 0.5M EDTA to a final concentration of 10mM. The ChIP was repeated eight times for each strain and then pooled, transferred to a mini dialysis tube (GE Healthcare Mini Dialysis Kit) and dialysed two times for 2 hours against 500ml of lysis buffer with reduced SDS (50mM HEPES-KOH pH 7.5, 150mM NaCl, 1% Triton X-100, 0.1% sodium deoxycholate, 0.05% SDS) before concentration by speed vacuum. The chromatin was then dissolved in western blot loading buffer and boiled at 95°C for 5 min before an aliquot was loaded on a 12% SDS polyacrylamide gel, transfer to a PVDF membrane and immunoblotting with anti-H3 antibody. Then a further Western analysis with anti-H3K4me1, anti-H3K4me2 and anti-H3 was performed using sample inputs equalized for the same amount of H3.

**ChIP-Seq Analysis**

ChIP-Seq reads were mapped to sacCer3 *S. cerevisiae* genome assembly using BWA and parallel version of pBWA (0.5.9-r32-MPI-patch2). After mapping, uniquely mapped were filtered (-q 1) and PCR duplicates (reads with same start and mapped to same strand) were removed using Samtools (rmdup) [11]. Read lengths were computationally extended to 150bp strand specifically and stored in Bam format. The Bam files were subject to coverage calculation using the bamCoverage utility of deepTools [12] for a sliding window of 40bp, normalized using RPKM parameters and stored in BigWig format. Additionally, these files were then subtracted by the mock IP control. Thus, all the resultant data were directly comparable because of RPKM normalization and control subtraction. Several data matrices were created for different loci of TSS +/-1.5KB, averaged metagene from the bigwig files via computeMatrix and visualized using plotHeatmap utilities of deepTools (also referred to as high resolution intensity plots or heatmaps). GO analysis was performed using the web-based tool FuncAssociate [13]. Composite profile plots were generated using custom code in R statistical language where the input was the above computed matrices using deepTools. BigWig files were visualised as coverage tracks using the UCSC genome browser [14].

For qChIP experiments, reads were first aligned to sacCer3 reference and then to sPombe reference genomes (obtained via PomBase; [15]). Note that reads were independently aligned to both genomes thus allowing for common mappings as compared to left-over read alignment to reduce the mismatch bias. Percentage of reads aligned per sample per genome was documented in a tabular format.

**RNA-Seq Analysis**

RNA-seq data were processed mostly using Tuxedo suite and custom scripts for further downstream analysis [16]. Tuxedo suite consists of TopHat (aligns the reads and map them to the genome) [17], Cufflinks (uses the read alignment map to assemble reads into transcripts) [18], Cuffdiff (takes aligned reads from multiple conditions and performs differential gene expression analysis) [19]. The counts are calculated and recorded in an expression unit of 'Fragments Per Kilobase of exon model per Million mapped reads' (RPKM). We used a cutoff of FPKM>4 for a gene to be called as expressed. Plots were generated using cummeRbund and custom scripts written in R statistical language.
